# Understanding Mechanisms of Chamber-Specific Differentiation Through Combination of Lineage Tracing and Single Cell Transcriptomics

**DOI:** 10.1101/2021.07.15.452540

**Authors:** David M. Gonzalez, Nadine Schrode, Tasneem Ebrahim, Kristin G. Beaumont, Robert Sebra, Nicole C. Dubois

**Affiliations:** Department of Cell, Developmental, and Regenerative Biology, Icahn School of Medicine at Mount Sinai, New York, NY 10029, USA; Mindich Child Health and Development Institute, Icahn School of Medicine at Mount Sinai, New York, NY 10029, USA; Black Family Stem Cell Institute, Icahn School of Medicine at Mount Sinai, New York, NY 10029, USA; Department of Genetics and Genomic Sciences, Icahn School of Medicine at Mount Sinai, New York, NY 10029, USA; Sema4, a Mount Sinai venture, Stamford CT, 06902

## Abstract

The specification and differentiation of atrial and ventricular myocardial cell types during development is incompletely understood. We have previously shown that *Foxa2* expression during gastrulation identifies a population of ventricular fated progenitors, allowing for labeling of these cells prior to the morphogenetic events that lead to chamber formation and acquisition of bona fide atrial or ventricular identity. In this study, we performed single cell RNA sequencing of *Foxa2Cre;mTmG* embryos at the cardiac crescent (E8.25), primitive heart tube (E8.75) and heart tube (E9.25) stage in order to understand the transcriptional mechanisms underlying formation of atrial and ventricular cell types at the earliest stages of cardiac development. We find that progression towards differentiated myocardial cell types occurs primarily based on heart field progenitor identity, and that different progenitor populations contribute to ventricular or atrial identity through separate differentiation mechanisms. We identified a number of candidate markers that define such differentiation processes, as well as differential regulation of metabolic processes that distinguish atrial and ventricular fated cells at the earliest stages of development. We further show that exogenous injection with retinoic acid during formation of the cardiac primordia causes defects in ventricular chamber size and is associated with dysregulation in FGF signaling in anterior second heart field cells and a shunt in differentiation towards orthogonal lineages. Retinoic acid also causes defects in cell-cycle exit in myocardial committed progenitors that result in formation of hypomorphic ventricles with decreased expression of important metabolic processes and sarcomere assembly. Collectively, our data identify, at a single cell level, distinct lineage trajectories during cardiac progenitor cell specification and differentiation, and the precise effects of manipulating cardiac progenitor field patterning via exogenous retinoic acid signaling.

## Introduction

The heart is one of the earliest organs to form and plays a key role in providing nutrients to the embryo at early stages of development. The morphogenesis events that characterize its formation are complex, due in part to the multiple progenitor populations that contribute progeny to distinct regions of the heart (Kelly et al., 2014). Clonal lineage tracing experiments have demonstrated that all cells in the heart are derived from a *Mesp1^+^* progenitor that originates from the cardiac mesoderm and ingresses into the lateral plate mesoderm (LPM) shortly after gastrulation (Chabab et al., 2016; Chiapparo et al., 2016; Lescroart et al., 2014). These *Mesp1+* cells then form two distinct progenitor populations termed the first heart field (FHF) and the second heart field (SHF), which together form the primordial heart structure known as the cardiac crescent (CC). The FHF forms the anterior region of the cardiac crescent consisting of a transient population that begins to express markers of differentiated cardiomyocytes such as *Nkx2-5* and *Tnnt2* before the initiation of primitive heart tube (PHT) formation, where the CC balloons out anteriorly and undergoes looping to form the primordial ventricle (Meilhac et al., 2014). The SHF forms posteriorly to the FHF at the CC stage and is characterized molecularly by expression of *Isl1*. In contrast to the FHF, the SHF persists as a progenitor population for longer, continuing to proliferate and provide the developing heart tube with differentiating cell types that migrate and become incorporated into the developing heart tube sequentially. More recent work has shown that the SHF can be divided into an anterior *Tbx1-*expressing domain and a posterior *Foxf1/Hoxb1-*expressing domain that contribute to the arterial and venous poles respectively (Bertrand et al., 2011; Rana et al., 2014; Roux et al., 2015). Derivatives from the FHF within the four chambered hearts are found primarily in the anterior structure of the left ventricle (LV) and to a lesser extent in the right ventricle, while the anterior second heart field (aSHF) contributes primarily to the right ventricle (RV) and outflow tract (OFT) with contribution to the atria from both the FHF and SHF (Meilhac et al., 2014; Zaffran et al., 2004). In this manner, regional contributions to the developing heart from heart field progenitors are influenced in large part by position, both in terms of anterior/posterior patterning within the heart itself as well as the location of the progenitor cells. Spatial differences in progenitor position are accompanied with differences in signaling environments the cells experience, with surrounding structures such as the developing gut tube providing important instructive cues that drive segregation and differentiation of these progenitors (Gavrilov and Lacy, 2013; Miquerol and Kelly, 2013).

The direct mechanistic consequences of the spatiotemporal distribution of cardiac progenitor cells in the early embryo remain incompletely understood, as detailed molecular studies of the heterogeneity of transiently occurring cells has only recently become possible with advancements in single cell sequencing technology. Such mechanistic studies in progenitor cells of the cardiovascular lineage are particularly important for the understanding of congenital heart defects (CHDs), many of which are thought to originate due to dysregulation of early heart development (Kloesel et al., 2016; Pierpont et al., 2018). CHDs often involve large structural defects in multiple regions of the heart that arise from the same progenitor, placing the heart field progenitors at the center of investigations into CHD disease etiology. Yet other developmental defects such as atrial/ventricular chamber size imbalances are not adequately described by the existing heart field model, as the FHF and SHF cells each contribute to both atrial and ventricular chambers. This raises the question of whether dysregulation of specific subpopulations within heart field progenitors, or dysregulation in specific processes along the atrial-ventricular differentiation trajectory might contribute to CHD formation.

Underlying these concepts remain a number of open questions about the precise mechanisms that govern atrial and ventricular specification and differentiation. The role of retinoic acid (RA) signaling as an external morphogen gradient driving atrial and ventricular differentiation is well-established and has been used to inform chamber specific differentiation protocols in pluripotent stem cell (PSC) models (Devalla et al., 2015; Lee et al., 2017; Perl and Waxman, 2020; Zhang et al., 2011). However, the transcriptional mechanisms downstream of this signaling pathway and its activity as a gradient, as well as the ordered cascade of genes that help to define acquisition of atrial or ventricular identity are not well understood. A number of efforts have attempted to understand these mechanisms by profiling the atrial and ventricular chambers separately and performing bulk and single cell RNA sequencing on the four-chambered heart as early as embryonic day 9.0 (E9.0) (DeLaughter et al., 2016; Li et al., 2016). These studies may however be limited in that they may miss key transcriptional processes that occur prior to the process of chamber morphogenesis and differentiation. More recent studies have profiled the transcriptome of the early developing heart at earlier stages shortly after gastrulation as the lateral plate mesoderm and cardiac crescent structures form (Ivanovitch et al., 2021; Lescroart et al., 2018; Tyser et al., 2021), or as the process of chamber morphogenesis is just beginning (Mantri et al., 2021; de Soysa et al., 2019). These studies helped to shed new light on the transcriptional heterogeneity of heart field progenitors at these stages and identified new markers that segregate these subtypes from one another in a spatiotemporal manner. They also identified transcriptional signatures for a number of other differentiated cell types at later stages and demonstrated how comparative analysis of scRNAseq data from WT and genetic knock-out models could shed light on transcriptional mechanisms underlying lineage relationships between heart field progenitors and their progeny. However, these studies did not explore in detail the specification and differentiation of atrial or ventricular cells particularly at early stages prior to acquisition of chamber-specific marker expression. We and others have recently expanded on a long-standing concept hypothesizing that atrial/ventricular lineage segregation may be occurring far earlier during mammalian development than previously understood (Bardot et al., 2017; Chabab et al., 2016; Garcia-Martinez and Schoenwolf, 1993; Keegan et al., 2004; Lescroart et al., 2014; Yutzey and Bader, 1995). We showed that transient expression of *Foxa2* in the primitive streak labels a population of early ingressing cardiac mesoderm which migrates at the leading edge of the LPM. *Foxa2* lineage-tracing experiments have shown that these early migrating mesodermal progenitors contribute to the majority of the ventricular, but not atrial myocardium (Bardot et al., 2017). More recent work has further corroborated this finding, demonstrating through single cell sequencing and genetic lineage tracing of *T* and *Foxa2* expressing cells that the ventricular and atrial cell types arise from spatially and molecularly distinct regions in the primitive streak (Ivanovitch et al., 2021). Thus, *Foxa2* lineage tracing may serve as a useful tool for labeling the ventricular progenitors prior to expression of chamber-specific markers.

Bulk RNA sequencing and differential expression analysis of *Foxa2* lineage-traced positive and negative cells within the cardiac mesoderm at E6.5 demonstrated upregulation of Notch pathway components within ventricular progenitors, indicating that early segregation and migration of these lineages may influence differences in signaling cues received by either population that in turn influences downstream mechanisms of differentiation (Bardot et al., 2020). However, it remains unclear whether atrial and ventricular fated progenitors exist as molecularly distinct subtypes within the first and second heart field, and whether atrial or ventricular fate is plastic at this time. The potential heterogeneity of these populations may also be masking sub-type specific differences in gene expression and signaling that govern atrial and ventricular differentiation from different heart field progenitors.

In order to interrogate the molecular identity of atrial and ventricular specific progenitors, as well as lineage relationships between progenitors and their differentiated progeny, we performed single-cell sequencing of dissected cardiac regions from *Foxa2Cre;mT*mG embryos at the cardiac crescent (E8.25), primitive heart tube (E8.75) and heart tube (E9.25) stages. Through expression of EGFP within the transcriptome, we were able to label ventricular progenitors within specific heart field populations without the need for cell sorting. RNA velocity measurements revealed relationships between distinct heart field progenitors and atrial/ventricular cells which were divided into various subtypes based on their transcriptome. Through differential expression analysis of atrial and ventricular progenitors we identified a number of candidate markers that correlate with atrial or ventricular differentiation over time. Lastly, we sought to understand how these transcriptomic differences are affected by exogenous RA signaling which has been known to modulate atrial and ventricular fate during early cardiac development. We find that exogenous RA negatively impacts ventricular development by preventing differentiation from separate progenitors towards a ventricular fate. This leads to defects in ventricular size and shifts in differentiation trajectories from progenitors towards other lineages.

## Results

### Generation of the transcriptomic landscape and identification of early cardiac ventricular lineage cells at single cell resolution

In order to label atrial and ventricular fated progenitors during the early phases of murine development prior to and during chamber formation, we crossed *Foxa2Cre* mice with *mTmG* mice to produce *Foxa2Cre;mTmg* embryos. Immunofluorescence analysis of *Foxa2Cre;mTmG* embryos at E14.5 after formation of the four chambered heart demonstrated robust EGFP labeling of the ventricular tissue, demonstrating that lineage tracing of *Foxa2* marks a population of ventricular fated progenitors (Bardot et al., 2017) (**Figure 1A**). We wished to profile the heterogeneity of cell types involved during early cardiac specification and differentiation and to capture the transcriptional events controlling morphogenesis of atrial and ventricular chambers. In order to understand these events and the distribution of ventricular fated progenitors within this landscape, we conducted single cell RNA sequencing (scRNAseq) on *Foxa2Cre;mTmG* embryos at the cardiac crescent (CC; E8.25), primitive heart tube (PHT; E8.75) and heart tube (HT; E9.25) stages (**Figure 1B**). To enrich for the populations of interest, we sub-dissected the cardiac regions from *Foxa2Cre;mTmG* embryos as well as the surrounding areas comprised of the developing gut tube and pharyngeal structures (**Supplementary Figure 1**) which are known to provide important signaling cues that govern cardiac differentiation and morphogenesis (Miquerol and Kelly, 2013). Three age-matched embryos per stage were pooled and a minimum of 10,000 cells sequenced at each stage (CC: 14,355, PHT: 13,036, HT: 21,318) targeting a depth of ∼50,000 reads/cell, resulting in an average of 2,000-4,000 genes called per cell.

**Figure 1.**
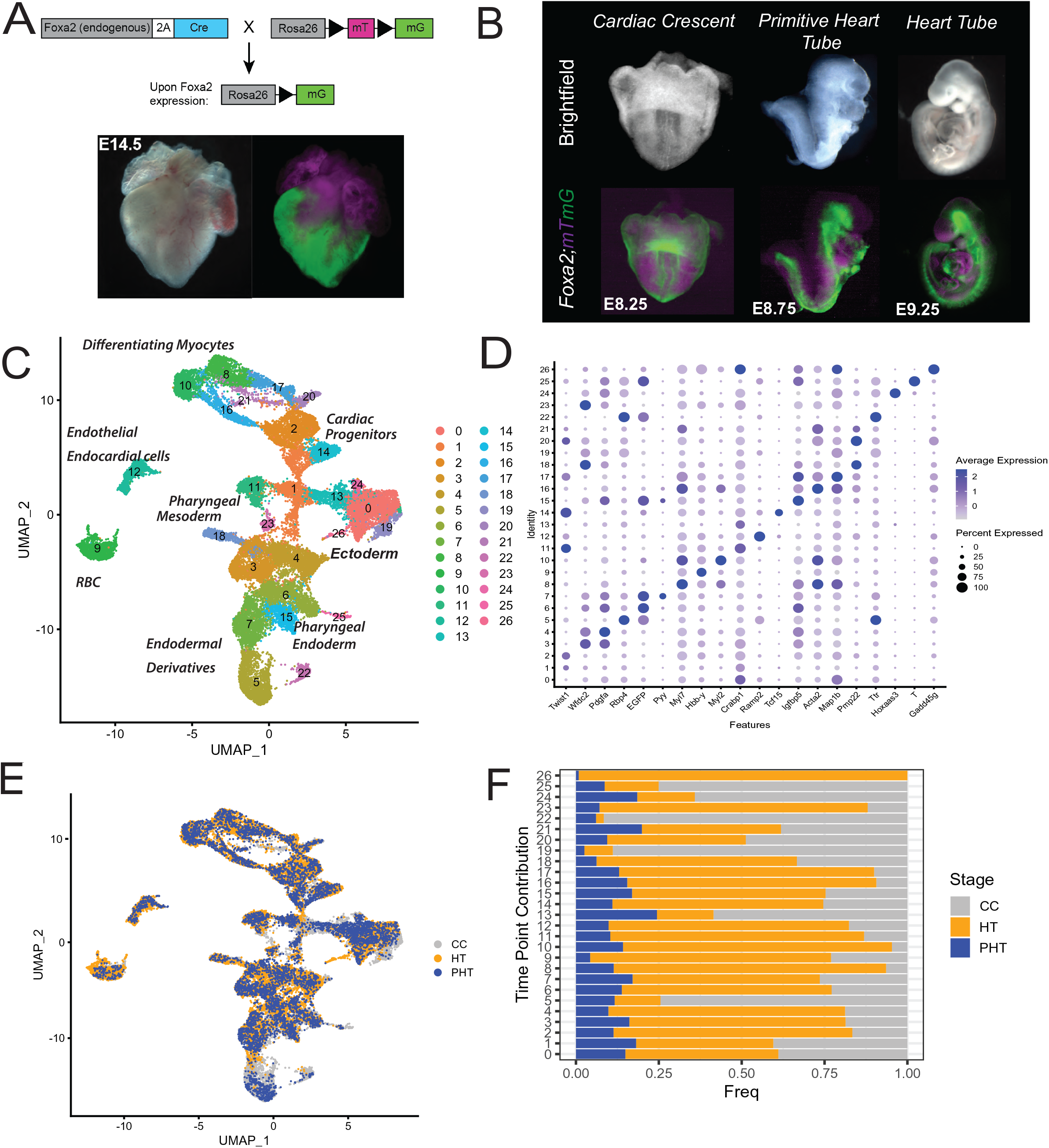
Single-cell Sequencing of *Foxa2* Lineage Traced Embryos Reveals a Changing Landscape of Cardiac Development and Surrounding Tissues. **A)** *Top*: Mating strategy to generate *Foxa2Cre;mTmG* mice. Crosses were done each time using *Fox2Cre* males crossed with *R26;mTmG* females. *Bottom*: Lineage tracing of Foxa2-mTmG mice shows enriched labeling of cells that form the ventricular but not atrial tissue. **B)** Representative images of embryonic stages sequenced, including the cardiac crescent (CC;8.25), primitive heart tube (PHT;8.75), heart tube (HT;9.25). **C)** UMAP clustering of all cells sequenced, demonstrating presence of differentiated myocardial cells, cardiac progenitors, pharyngeal cell types, endodermal derivatives, and endothelial/endocardial cells. **D)** Bubble plot of top differentially expressed gene for each cluster. Differentially expressed genes for each cluster were calculated using a Wilcoxon rank-sum test with p-value <0.01. **E)** UMAP of all cells sequenced showing contribution from samples from each stage: *grey =* CC; *blue =* PHT; *orange =* HT. **F)** Quantification of overall contribution of cells from any one particular embryonic time point. Frequency is calculated as the number of cells in a given cluster relative to the total number of cells in the sample.

We performed clustering and differential gene expression analysis at each stage to identify several distinct populations of mesodermal and early cardiac populations, as well as differentiating cardiomyocytes (**Supplementary Figure 2A-C**). Additionally, we identified several populations of definitive ectoderm and endoderm-derived cell types, including neural crest cells that contribute to both the heart and head. Importantly, we find that we can label *Foxa2* lineage-traced cells through identification of *EGFP+* cells within our transcriptomic data set, without the need for cell-sorting that might have resulted in loss of rare cell types. *Foxa2* is prominently expressed in endoderm and endoderm-derived lineages. Accordingly, we identified populations of *EGFP+* cells within endoderm populations (*Epcam+, Foxa2+, Sox17+*). Foxa2 lineage-traced cells were also present within mesoderm populations, where they label a population of ventricular fated cardiomyocytes. We wished to confirm that expression of *EGFP* correlated with expression of markers of ventricular identity at the PHT and HT stages, when ventricular cells become physically separated from their atrial counterparts and take on bona-fide ventricular transcriptional identity through expression of markers such as *Myl2* and *Irx4.* As expected *EGFP+* cells were found within *Irx4+* cells at the HT and PHT stage but were largely absent within *Kcna5+* atrial cell types at the same stages **(Supplementary Figure 2B-C**). We also observed expression of *EGFP* within mesoderm cells that expressed early markers of the cardiac lineage such as *Pdgfra* and *Nkx2-5,* indicating that our strategy allows us to label ventricular fated cells within early progenitors prior to expression of chamber-specific genes.

We next merged the data from the three time points in order to create a time-resolved map of the developing heart and surrounding tissues (**Figure 1C**). We identified a total of 26 clusters that separated broadly according to germ layer identity. We performed differential expression analysis on a cluster-by-cluster basis to determine gene signatures that define the individual populations (**Figure 1D**). This analysis identified differentiated cardiomyocytes and associated progenitors, pharyngeal endoderm and mesodermal cell types, as well various definitive ectoderm and endodermal cell types among others. In order to determine the effects of embryonic time on cluster identity, we calculated the frequency of contribution to each cluster across the CC, PHT and HT samples (**Figure 1E/F**). We found that more differentiated clusters comprised of *Tnnt2+* cardiomyocytes (Clusters 8 & 10) were primarily composed of cells from the PHT and HT embryo samples, while cardiac progenitors of a cluster expressing high levels of the primitive streak marker *T* had higher contributions from CC stage samples (Cluster 25). In summary, combining single cell technology with Foxa2 lineage tracing allowed us to generate a map of the transcriptomic landscape across real developmental time of the cardiac lineage and to identity ventricular fated cells at various time points along their differentiation trajectory.

### Clustering of cardiovascular cells reveals lineage relationships across time between heart field progenitors and differentiated progeny

Previous molecular and fate mapping studies have provided important insight into lineage relationships between heart field progenitors and the cardiomyocytes that make up the chambers of the heart (Francou and Kelly, 2016; Kelly et al., 2014; Lescroart et al., 2014). However, it remains largely unclear if differences between heart field progenitors such as the FHF and aSHF/pSHF represent fundamentally distinct transcriptional programs with differing lineage potential, or if differences in their marker expression is a consequence of spatiotemporal differences in the embryo during sequential events of cardiogenesis. More recent studies have profiled these populations with increasing resolution, and demonstrated increased molecular heterogeneity of populations at early stages (Ivanovitch et al., 2021; de Soysa et al., 2019; Tyser et al., 2021). However, it remains to be shown whether these subpopulations represent lineage restricted progenitors or whether these subtypes are intermediate cell-states in the process of differentiation.

In order to focus on the heterogeneity and lineage trajectories of the cardiac lineages, we subset the differentiated cardiomyocytes and their progenitors along with neural crest, endocardial and endothelial cells, as well as epicardial cells at each stage (CC, PHT, HT) and reclustered these cells **(Supplemental Figure 3A-C).** This allowed us to create a snapshot of the developing cardiac lineages at discrete stages of embryonic development. To understand dynamics of how these cell types change throughout embryogenesis, we merged the CC, PHT and HT samples to generate a combined data set of 26 distinct clusters resolving different heart field progenitors, early differentiating cardiomyocytes of mixed chamber identity, as well as differentiated cardiomyocytes that segregate into atrial or ventricular identity (**Figure 2A**). Progenitor cell types and differentiated cells segregated from one another in part based on cell cycle state, an important covariate in scRNASeq data that we did not regress in our data due to its biological importance in rapidly developing tissues **(Figure 2B).** We performed differential gene expression analysis across all 26 clusters to identify candidate markers that defined each population **(Figure 2E)**. We identified multiple *Pdgfra+* early mesoderm populations that clustered separately from more differentiated *Nkx2-5+/Tnnt2+* cardiomyocytes, and found that these progenitors were more commonly found in the S phase of the cell cycle compared to their differentiated progeny, which were more commonly found in G2M/G1 phase **(Figure 2B/C)**. These data are consistent with the role of cardiac progenitors, particularly the SHF, as a stable pool of progenitors that proliferates to maintain the progenitor pool and contributes differentiating cell types to the growing heart (Meilhac et al., 2014; Miquerol and Kelly, 2013). *Pdgfra+* progenitor clusters separated into somitic mesoderm, LPM or SHF identity based on expression of *Fst, Pmp22* and *Isl1* respectively (**Figure 2C**). The *Isl1^+^* SHF progenitors were further subdivided into anterior and posterior domains, both of which co-expressed *Nkx2-5* while maintaining proliferative capacity as indicated by the predominance of cells in S phase within this cluster. The aSHF population was identified through expression of *Tbx1*, while the pSHF clustered separately and expressed higher levels of *Aldh1a2* and *Hoxb1*.

**Figure 2.**
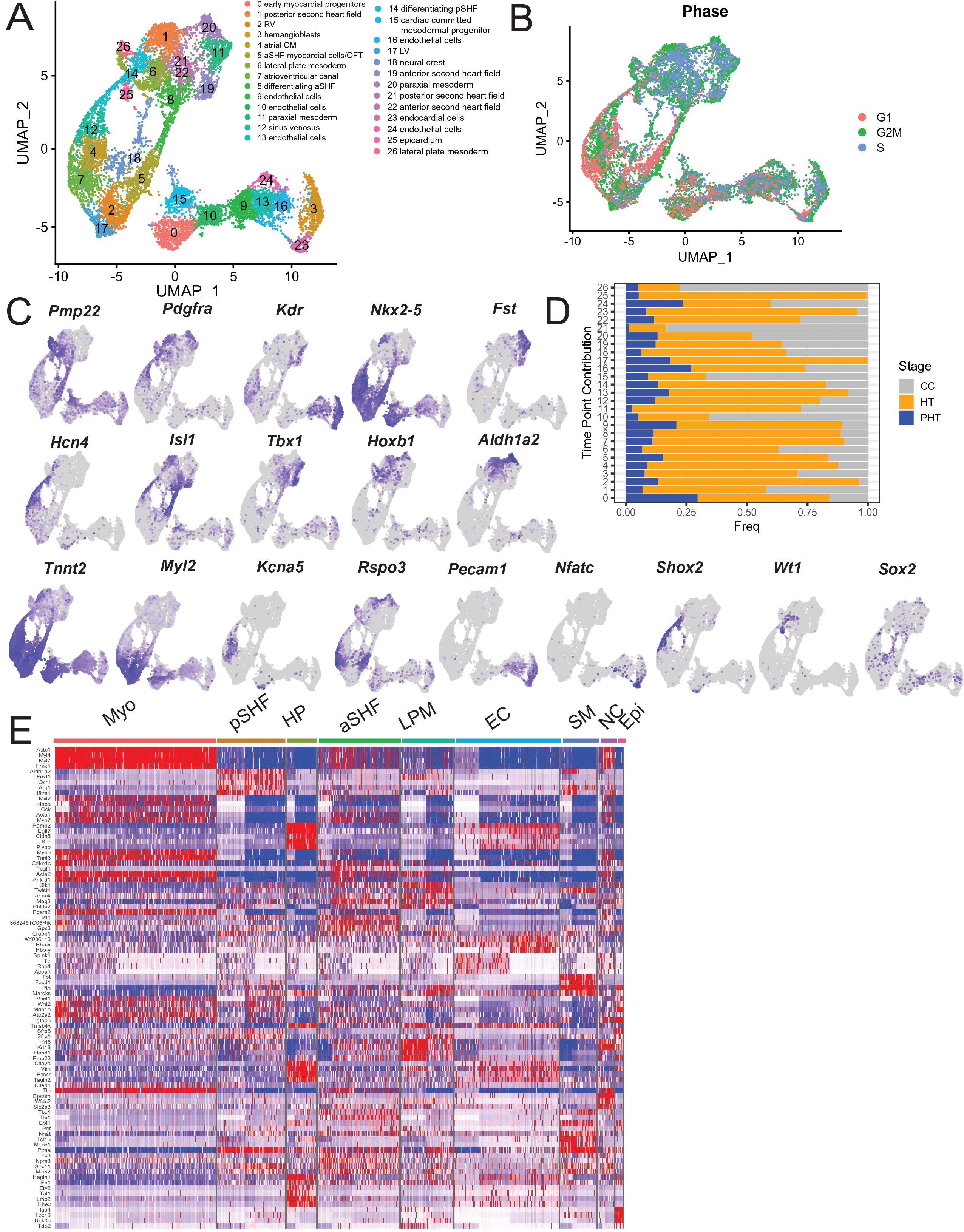
Sub-clustering of Cardiac Regions Reveals Heterogeneity of Progenitor and Differentiated Cell Types. **A)** UMAP clustering of cardiac regions, demonstrating presence of multiple progenitor and differentiated cell populations. **B)** Cell cycle phase scoring for each cell projected onto UMAP embedding. **C)** FeaturePlots of key markers for progenitor cell types, differentiated myocardial cells as well as other cell types including epicardium, neural crest, and endothelial cells. **D)** Contribution of cells from each time point to individual clusters (*grey =* CC; *blue =* PHT; *orange =* HT). Frequency is calculated as the number of cells in a given cluster from one time point relative to the total number of cells in the sample**. E)** Heatmap demonstrating top 5 most differentially expressed genes for broad categories of cell types. CM = cardiomyocytes; pSHF = posterior second heart field; HP = hemangioblast progenitor; aSHF = anterior second heart field; LPM = lateral plate mesoderm; EC = endothelial/endocardial cell; NC = neural crest; Epi = epicardium.

We did not observe a specific population of FHF progenitors in our merged data set, but did observe that the CC sample contributed a group of *Tnnt2+* cells that co-expressed *Hcn4* and *Tbx5* to our merged object **(Supplementary Figure 4A)**, and that these cells could be found within multiple differentiating cardiomyocyte clusters (Clusters 7,12 & 14). The clustering of the FHF cell types within the CC together with differentiating cardiomyocytes underscores the identity of the FHF as a transient progenitor population that, despite its close spatial association with the SHF in the early cardiac crescent, transcriptionally more closely resembles differentiating cardiomyocytes than a heart field progenitor. It is likely that shared expression of markers such as *Tbx5* and *Hcn4* within the FHF cells and expression of these markers in the sinus venosus and other differentiating cells from the pSHF may be driving clustering of these cell types.

Differentiated cardiomyocytes expressing *Tnnt2* and *Nkx2-5* segregated into several subpopulations including atrial and ventricular cells that expressed *Kcna5,* and *Myl2* respectively. Of note, *Myl7* expression, a classic marker of atrial identity in adult cardiomyocytes was expressed in the majority of *Tnnt2+* cells, indicating that canonical post-natal atrial/ventricular markers may be expressed in different patterns and perform different functions during early development **(Supplementary Figure 4B).** *Myl2+* ventricular cardiomyocytes segregated into two clusters representing left and right ventricular cells characterized by expression of *Hand1/Pln* and *Cited1/Cck,* respectively **(Supplementary Figure 4C)**. We also observed several populations (clusters 0 & 5) that expressed a combination of atrial and ventricular markers **(Figure 2C).** We hypothesized that these represent newly differentiated cardiomyocytes that cluster due to a shared origin rather than atrial/ventricular identity. Accordingly, cluster 0 appeared to cluster closely together with a *Pmp22+* population which segregated away from the SHF and LPM progenitors (cluster 15). This cluster had increased expression of *Nkx2-5* compared to the LPM, perhaps indicating that cluster 15 represented a separate group of mesoderm progenitors that has already committed toward a cardiac fate.

In addition to differentiating cardiomyocytes, we identified a population of *Sox2-*expressing neural crest cells, as well as a population of *Wt1/Tbx18*-expressing epicardial cells (**Figure 2C/E**) which arises from a progenitor population known as the pro-epicardial organ (PEO) that develops separately from the FHF and SHF (Niderla-BieliŃska et al., 2019). We also identified populations corresponding to the *Shox2+/Hcn4+* sinus venosus (SV) and *Rspo3+* atrioventricular canal (AVC). Both of these clusters were found in close proximity to the atrial cardiomyocytes, consistent with their shared occupancy of the venous pole within the developing heart tube as the atrial and ventricular chambers begin to form. Of note, we also identified a separate *Rspo3+* population (cluster 5) that clustered separately from the atrioventricular canal (AVC) cells (cluster 7), indicating that *Rspo3* may not be a specific maker for this lineage at all stages of development. R-spondins have been shown previously to amplify sensitivity to Wnt ligands (Lebensohn and Rohatgi, 2018). Given the multiple cycles of Wnt activation and inhibition at different stages of cardiac development (Gessert and Kühl, 2010), we hypothesized that expression of *Rspo3* in these cells may correlate with canonical Wnt activation driving proliferation of newly differentiated cardiomyocytes, though further studies would be needed to determine this in detail.

In addition to the myocardial cell types we identified several populations of *Pecam1+* endothelial cells (clusters 9,13,16), which clustered closely together with a population of *Nfatc1+* endocardial cells (cluster 23). Of note, a number of *Pecam1+* cells also co-expressed *Nkx2-5,* which may represent a population of angioblasts that have been reported to contribute to the hemogenic endothelium outside the cardiac crescent (Zamir et al 2017.). Both populations clustered closely to a *Kdr+* progenitor population (cluster 3) that may represent a shared hemogenic/endocardial/endothelial progenitor, and were also closely associated with the previously described *Pmp22+/Nkx2-5+* mesoderm progenitors. Given the separation of this population from the LPM and toward the endothelial/endocardial cell types it is possible that this may represent a bipotent mesodermal progenitor contributing to myocardial and endothelial/endocardial lineages, though further studies would be needed to understand this population in more depth.

In an effort to understand how the landscape of cardiac development changes over real developmental time, we quantified the contribution from each of the three developmental stages (CC, PHT, HT) to each of the identified cell populations (clusters 1-26). This identified a number of clusters that were overrepresented at specific embryonic time points. Early progenitors such as the LPM were comprised mostly of cells from the CC stage (clusters 6 & 26), while more differentiated cardiomyocytes that had acquired atrial or ventricular identity (clusters 2,4 & 17) were found to arise primarily from samples at the PHT and HT stage (**Figure 2D**). The *Isl1+* SHF progenitors in contrast had a more balanced contribution from all three stages, in accordance with their persistence at the arterial and venous poles throughout the morphogenetic processes of heart tube formation, looping, and initiation of chamber formation. *Wt1+* epicardial cells were derived primarily from the E9.25 HT sample with small contribution from the E8.75 PHT stage, consistent with the later migration of epicardial cells from the POE which initially develops distally from the heart.

Collectively, we have generated a map of early cardiac development that recapitulates the transcriptional landscape at the CC, PHT and HT stages and the changes that occur across these stages at single cell resolution. We identified several populations belonging to the LPM and aSHF/pSHF progenitors, as well as a molecularly distinct *Pmp22+* LPM population with increased *Nkx2-5* expression that clusters separately from the LPM and heart field progenitors. By quantifying the contribution of time-resolved samples to our merged data, we captured the disappearance of early progenitors between early and late-stage samples, concomitant with an emergence of more differentiated cell types that express the canonical atrial/ventricular markers, or a combination of both. Interestingly, we find that despite their position adjacent to the SHF within the CC, when compared to myocardial cell types across multiple stages of development, *Hcn4+/Tbx5+* cells from the CC stage (representing FHF cells) more closely resemble differentiating cardiomyocytes. This underscores the growing understanding of the FHF as a transient differentiating population rather than a true progenitor counterpart to the SHF population within the developing heart.

### Identification of signaling pathways and processes underlying cardiomyocyte heterogeneity in the early embryo

We were intrigued to find in our data the presence of multiple differentiated *Tnnt2+* cardiomyocyte clusters, only some of which could be identified purely on the basis of atrial/ventricular or left/right identity. We hypothesized that these might represent different functional subtypes of cardiomyocytes, or various intermediary stages of differentiation. In order to more comprehensively understand differences between these cardiomyocyte subtypes, we performed differential expression analysis on a cluster-by-cluster basis to identity the top differentially expressed genes in each subtype (**Figure 3A**). We then performed gene set enrichment analysis to identify differentially regulated processes in each cluster. Subsequent GO term analysis indicated that the populations separated into distinct clusters on the basis of chamber identity, differentiation status, and metabolic changes associated with increased contractile ability (**Figure 3B**).

**Figure 3.**
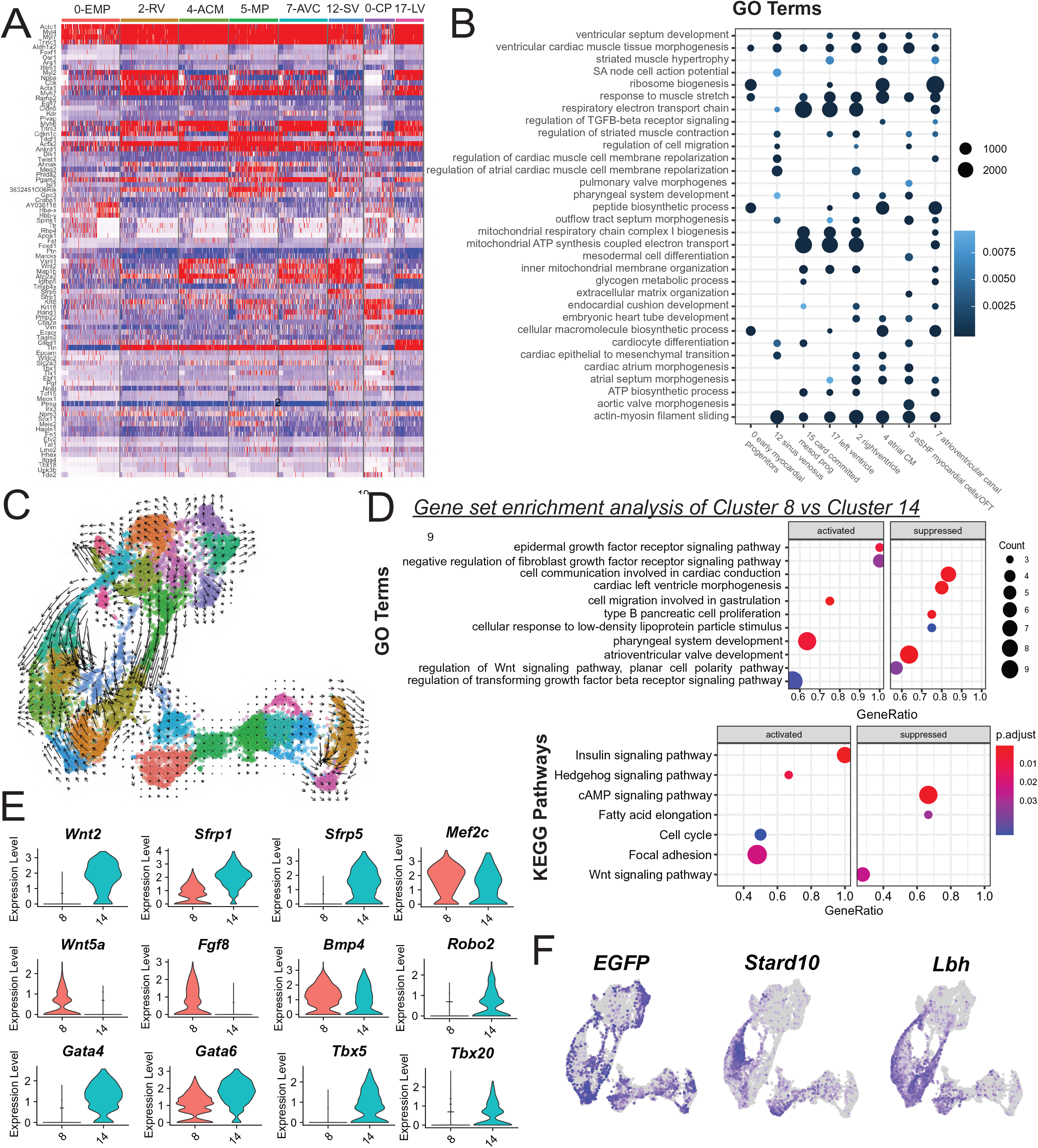
Characterization of Functional Cardiomyocyte Subtypes and Corresponding Lineage Relationships to Progenitor Cell Types. **A)** Heatmap demonstrating top 5 most differentially expressed genes for cardiomyocytes. **B)** Gene ontology analysis of differentially regulated processes in each cell type. **C)** RNA velocity demonstrating differentiation dynamics within cardiac subsets. **D)** GO term (top) and KEGG pathway (bottom) analysis of upregulated and downregulated processes between aSHF differentiating cells (cluster 8) and pSHF differentiating cells (cluster 14). **E)** Violin Plots of selected differentially expressed genes along differentiation trajectories. **F)** Expression of atrial/ventricular markers across UMAP clustering.

Not surprisingly, a number of GO terms were shared across several clusters, such as processes involved in “actin-myosin filament sliding”, consistent with a shared cardiomyocyte identity across the clusters analyzed. Others, such as “SA nodal action potential” were restricted to particular clusters such as the *Hcn4+* sinus venosus. We found that processes involved in mitochondrial ATP synthesis and other energetic processes seemed to segregate clusters into two broad categories of more/less mature cardiomyocytes, which require more energy as they become more contractile. These more mature cardiomyocytes also demonstrated increased expression of *Nppa,* indicating that they have begun to upregulate the chamber myocardium program that drives ballooning morphogenesis and encounter increased load within the chamber. Consistent with this finding, these cells expressed genes involved in processes of “ventricular and/or atrial differentiation” and “morphogenesis” again indicating an advanced maturation stage and acquisition of chamber-specific identity and transcriptional segregation from one another.

Clusters that did not show enrichment in energetic and morphogenesis processes instead were characterized by processes such as “ribosome biogenesis” and “peptide biosynthesis”. This led us to conclude that these clusters represented early cardiomyocytes and/or progenitors, many of which did not appear to have chamber specific identity yet, including our previously described *Pmp22+/Nkx2-5+* cardiac-committed mesoderm population (cluster 15) and another adjacent population of *Tnnt2+* cardiomyocytes that expressed markers of both atrial and ventricular cells in the four chambered heart. Both clusters had decreased expression of sarcomeric genes compared to more mature cardiomyocyte clusters. This suggested to us that these cells clustered together due to a shared and comparatively less mature differentiation state compared to the *Nppa+* expressing cardiomyocytes and that the strength of association along a shared developmental age or origin may predominate over emerging atrial and ventricular identity early during development.

A population we identified earlier as the AVC was characterized by processes such as “atrial septum morphogenesis” and “endocardial cushion development” (Cluster 7). Cells within this cluster expressed *Rspo3* which has been identified in other studies as a marker of the AVC. However, the expression of *Rspo3* was not exclusive to this cluster in our data, and also marked a population of *Tnnt2+* cells of unknown identity that clustered separately from bona-fide atrial or ventricular cells (Cluster 5). GO term analysis showed enrichment of processes involved in “aortic and pulmonary valve development”, yet cells within this cluster exhibited expression of sarcomeric proteins and processes involved in regulation of striated muscle contraction. These cells also clustered closely together to the *Isl1/Tbx1* expressing aSHF, leading us to hypothesize that this cluster might represent a mixed population of cell types derived from the aSHF which is known to give rise to cells of the RV and the outflow tract. In order to test this hypothesis, we re-clustered the cells within this population alone and found that they separated into three distinct clusters. These consisted of two population of *Myl2*+ cells as well as a separate population enriched for *Crabp1, Rgs5, Flna,* and *3632451O06Rik,* markers of the OFT (**Supplementary Figure 5**). This led us to conclude that cluster 5 represents a mixed population of cells arising from a common aSHF progenitor, further suggesting that the present cell clustering contains additional heterogeneity of cell types to be explored further.

In conclusion, we identified the existence of several cardiomyocyte subpopulations which were distinguished from one another primarily based on their differentiation state. We were intrigued to find that several populations representing intermediary differentiation states characterized by a less mature energetic phenotype were composed of mixed populations of cells. The propensity of these cells to cluster together despite expression of markers of different lineages, as evidenced by the existence of both OFT and myocardial cells clustering together in cluster 5, indicated that at early stages of differentiation transcriptional similarity due to a shared origin may predominate over cell-type specific gene expression.

### Characterizing differences in signaling and lineage relationship between heart field progenitors

Having hypothesized that the transcriptome of intermediary cell types seems at least in part defined by progenitor origin, we wished to better understand the lineage relationships between cardiac progenitors and their progeny. This question was further motivated by the presence of transcriptionally distinct progenitors that cluster far apart from one another, but seem to give rise to cells expressing similar genes. While clonal lineage tracing studies have provided important information about the eventual fate of various heart field progenitors, a comprehensive understanding of the molecular players involved in differentiation on a single-cell level is still needed. These studies may help to characterize dynamic changes in genes along differentiation trajectories, and answer long-standing questions about whether differentiation of FHF and aSHF/pSHF progenitors toward their progeny is performed through entirely distinct mechanisms.

We applied velocytoR to estimate RNA velocity across our cell types and to visualize differentiation dynamics and relationships between progenitors and their progeny. This analysis revealed several trajectories of differentiation, with cardiomyocytes that appeared to originate from separate regions of the heart field progenitors **(Figure 3C)**. The aSHF and pSHF populations gave rise to two parallel differentiation streams that contributed to differentiated cardiomyocytes in separate ways. Within this differentiation stream, clusters 8 and 14 appeared to capture rapidly differentiating cells exiting from the aSHF and pSHF respectively, as shown by the large unidirectional arrows in each cluster. Clusters comprising the atrial cells and the sinus venosus originated from the pSHF, as expected given its clonal lineage relationship to the venous pole of the developing heart tube. RNA velocity revealed two diverging fates of the aSHF toward the somitic mesoderm and cardiac lineages. Derivatives from the aSHF (e.g., RV and OFT) gave rise to populations of more differentiated ventricular cells, but also associate with *Sox2* expressing neural crest cell types that differentiated separately. These observations are consistent with important interactions between the neural crest lineage and the OFT that help drive septation of the aortic and pulmonary arteries during development. Accordingly, RNA velocity also identified a stream of differentiation within the neural crest cells whose terminal position diverged and rested with the AV, as well as cluster 5 which represented a mix of ventricular and OFT cell types derived from the aSHF. This association is consistent with the role of the NC in driving endocardial cushion development and the formation of the atrioventricular valves later during development. Interestingly, cells forming the AVC appeared to receive input from cell types arising from the pSHF derivatives that undertook a separate pathway of differentiation that diverged from cell types contributing to the *Kcna5/Nppa* expressing atrial chambers.

Based on this analysis, we found that ventricular cells arise from multiple sources, including the aSHF which gives rise to *Myl2* expressing cells in cluster 5, as well as an early population of myocardial progenitors derived from cardiac committed mesoderm cells (cluster 15). RNA velocity indicated that these mesodermal progenitors give rise to early myocardial cells of mixed atrial/ventricular fate in cluster 0, but may also contribute to cells within the endocardial/endothelial populations, perhaps indicating a bipotent ability that differentiates this mesodermal population further from other *Pmp22*-expressing LPM cells. Further studies will be necessary to understand if this population represents a separate pool of progenitors that contributes differently to the atrial/ventricular lineage, or if its transcriptional identity is driven by other mechanisms.

Given that our RNA velocity estimations appeared to capture differentiation events of the aSHF and pSHF, we next interrogated the underlying processes and signaling that drive this event across both heart fields. We performed differential gene expression between cluster 14 and 8, which appeared to represent rapidly differentiating cells arising from the pSHF and aSHF respectively (**Figure 3C**). We found that processes involved in “atrial and atrioventricular valve development” were downregulated in differentiating aSHF cells compared to differentiating pSHF cells, while processes involved in “pharyngeal system development and migration” were upregulated, consistent with the eventual progeny of these cell types. We identified differences in signaling of various pathways including Wnt, Hh, TGF-β and FGF signaling (**Figure 3D**). We plotted several differentially expressed candidate genes across both cluster and found that while differentiation of pSHF cells appeared to involve expression of canonical Wnt players such as *Wnt2*, *Sfrp1* and *Sfrp5*, the noncanonical Wnt ligand *Wnt5a* was highly expressed in differentiating aSHF cell types (**Figure 3E**). We found that differentiation of aSHF progenitors involved upregulation of *Bmp4* expression, as well as *Fgf8* which was expressed specifically in the differentiating aSHF, consistent with previously published data (De Zoysa et al., 2020; High et al., 2009; Pradhan et al., 2017). Differential gene expression across these two populations also showed differences in expression of key components of the cardiac gene regulatory network, such as shared expression of *Mef2c* while the pSHF differentiation stream appeared to demonstrate increased expression of *Tbx5* and *Tbx20*, as well as *Gata4* in comparison to differentiation of the aSHF. In this way, differentiation of aSHF and pSHF appears to occur through activation of discrete pathways deployed separately to drive differentiation along a cardiomyocyte lineage.

Lastly, we noticed that processes involved in “left ventricle development” were differentially regulated in cluster 14 **(Figure 3D).** We interrogated the expression of *Stard10* and *Lbh*, two markers of early atrial/ventricular identity across each differentiation streams and found that while *Stard10* expression was found exclusively within cells arising from cluster 14, *Lbh* expression could be found across both differentiation streams (**Figure 3F**). When investigating if ventricular fated *Foxa2* lineage traced cells are present within these populations, we found *EGFP+* cells to form a distinct line that segregated from the atrial cell types and appeared to extend from the LPM population to the *Myl2+* ventricular clusters. This expression pattern is reminiscent of the distribution of *Tbx5+/Hcn4+* cells from the CC that we previously identified as belonging to the FHF which is known to contribute to the LV during development **(Supplementary Figure 4).** These data indicate that the stream of differentiation captured by cluster 14 does not simply describe a population of differentiating atrial cells derived from the pSHF. Rather, its identity may represent a mix of pSHF derivatives as well as ventricular-fated cells from the LPM that differentiates through similar mechanisms as those employed by the pSHF in its differentiation towards atrial/SV cell types. This would suggest that intermediary cells of atrial/ventricular fate are defined moreso by their progenitor of origin and its transcriptomic signature during differentiation rather than eventual atrial/ventricular identity.

### Identification of markers of atrial/ventricular specification and differentiation across multiple differentiation states

Our previous data indicate that atrial and ventricular cells arise from multiple progenitor sources (e.g. SHF progenitors vs LPM subtypes), some of which may represent developmental intermediates at different stages. We wished to take advantage of this to identify and stratify markers of atrial/ventricular differentiation that are specific to atrial/ventricular cells at particular stages of differentiation. We began by performing pairwise differential expression analysis of atrial and ventricular chamber cardiomyocytes within the most differentiated cells in our data set (cluster 4 and clusters 2&17 respectively) (**Figure 4A/C**). We identified 3,695 differentially expressed genes, including known chamber-specific markers previously identified in the four chambered heart such as *Myl2, Mest,* and *Stard10* as well as a number of markers not previously associated with atrial/ventricular identity. We also identified other markers such as *Vsnl1* that label an early population of atrial cells in the heart tube shortly after looping but is expressed in all chambers in the adult (Ola et al., 2012). *Vsnl1* has also recently been shown to be expressed as early as the cardiac crescent in an anatomically distinct subtype of cardiac progenitors with FHF-like properties (Tyser et al., 2021), and thus may represent a subtype within the FHF which contributes specifically to the atria. Among our candidates enriched in ventricular cells was EGFP, further validating the use of the EGFP labeling within the transcriptome to enrich for ventricular cells. In order to predict potential functional consequences of these candidate genes, we performed GO and KEGG pathway enrichment analysis (**Figure 4D/E**). We observed upregulation of processes involved in regulation of calcium ion transport and mitochondrial respiratory chain complex assembly, and cardiac myofibril assembly within ventricular cells, consistent with the increase load on ventricular cells compared to their atrial counterparts. We also observed differences in a number of pathways including Hedgehog signaling, JAK/STAT and Ras signaling, as well as a number of metabolic processes such as thyroid hormone synthesis and steroid biosynthesis.

**Figure 4.**
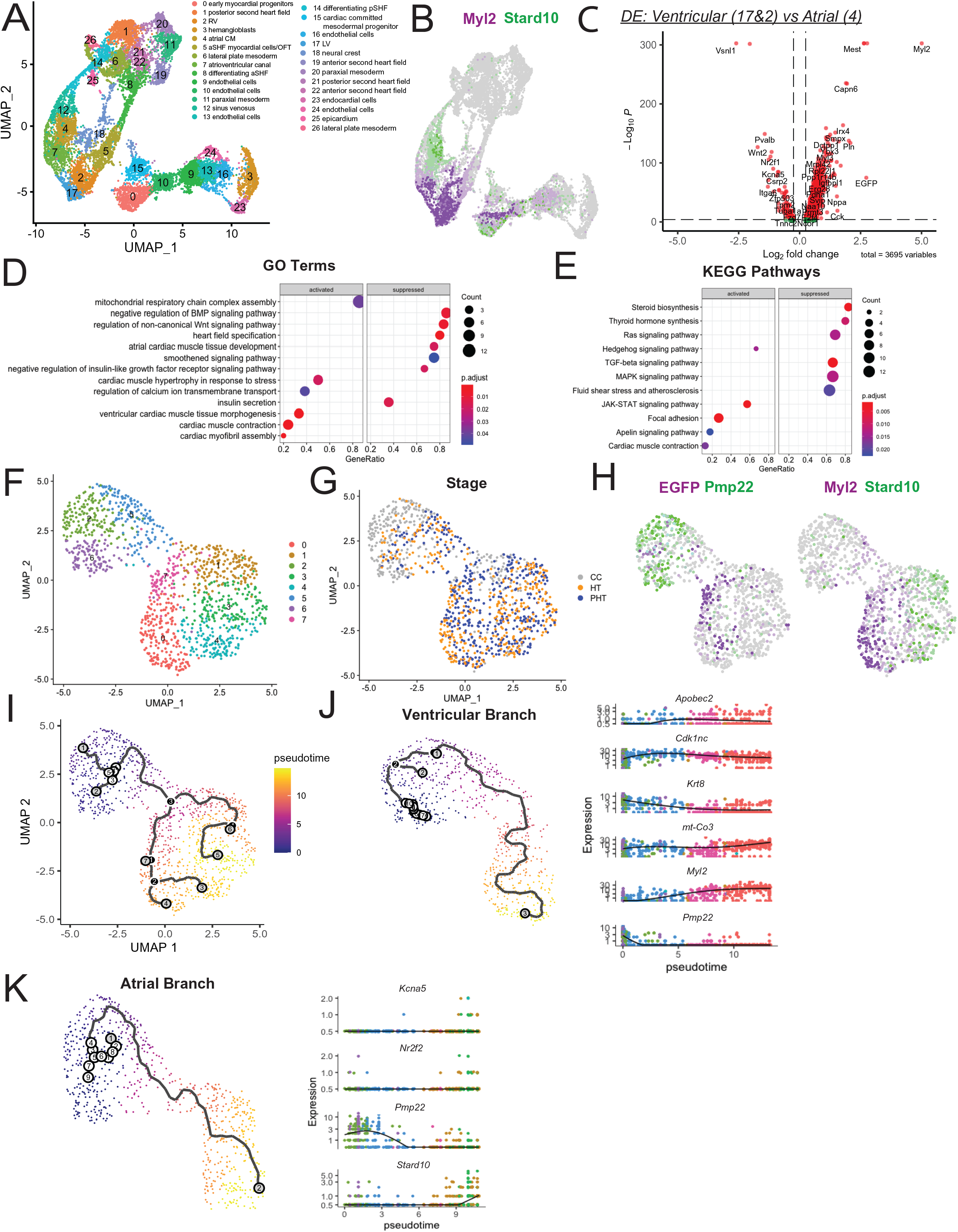
Differential Expression of Atrial and Ventricular Cells at Multiple Developmental Stages. **A)** UMAP clustering of cardiac subset. **B)** Feature plot showing segregation of atrial and ventricular cells into separate clusters (4 vs 2&17) as well as mixed contribution of atrial and ventricular cells to Cluster 0. **C)** Volcano plot demonstrating differential expression of ventricular (clusters 17&2) vs atrial cells (cluster 4). Differential expression analysis was performed using negative binomial distribution test in Seurat with p<0.01. **D)** Selected GO terms upregulated/downregulated in ventricular cells compared to atrial cells. **E)** Selected KEGG pathways upregulated/downregulated in ventricular cells compared to atrial cells. **F)** UMAP plot of cluster 0 and 15 following re-clustering. **G)** Time point contribution for individual samples. **H)** Feature plot showing expression of *EGFP* ventricular fated cells within *Pmp22+* mesodermal progenitor population and segregation of *Stard10+* atrial and *Myl2+* ventricular clusters. **I)** Pseudotime trajectory constructed in monocle3 demonstrating segregation into ventricular and atrial branches. **J)** Subset of ventricular branch with genes of interest plotted along pseudotime axis. **K)** Subset of atrial branch with atrial markers plotted along pseudotime axis.

In addition to bona fide clusters of atrial and ventricular identity in clusters 4, 2 and 17, we observed mixed expression of *Myl2* and *Stard10* in cluster 0 **(Figure 4B).** Based on the RNA velocity analysis, myocardial cells in this cluster appeared to be formed from a separate group of cardiac committed mesoderm cells derived primarily from cells at the CC stage (cluster 15). As indicated previously, both these populations were enriched for biosynthetic processes and had decreased expression of sarcomeric proteins, leading us to hypothesize that cells in cluster 0 with chamber-specific gene expression might represent a population of early differentiating cells that only recently acquired atrial/ventricular identity. We subset these populations and re-clustered them, finding that these formed separate groups of atrial and ventricular clusters that appeared from a common progenitor pool expressing the LPM marker *Pmp22* **(Figure 4H)**. We utilized Monocle3 in order to learn and visualize lineage trajectories within these cells (**Figure 4I**). We found that progenitor cells separated into atrial and ventricular lineages, allowing us to subset these lineages and plot expression of key markers along the pseudotime vector. We found that downregulation of LPM marker *Pmp22* within the ventricular lineage was soon followed by upregulation of cell cycle components (**Figure 4J**). In particular, we found that expression of apolipoproteins such as *Apobec2* were among the first genes upregulated along the differentiation trajectory towards the ventricular lineage, preceding the expression of canonical ventricular markers such as *Myl2* (**Figure 4K**). In contrast, expression of atrial markers *Stard10, Kcna5 and Nr2f2* appeared closely following downregulation of Pmp22, with fewer intermediate genes preceding the expression of canonical markers.

In summary, we identified a number of known and new candidate genes of atrial/ventricular identity within differentiated cardiomyocytes expressing chamber-specific markers. Experimental validation of the expression pattern and loss-of-function experiments of such candidates will be explored in future work to determine their spatiotemporal distribution within the developing heart and their functional relevance for early heart specification. We additionally identified a separate group of atrial and ventricular cardiomyocytes that appear to be derived from a separate progenitor source, and utilized the diverging trajectory of this progenitor to identify candidate genes that are expressed early during differentiation toward a ventricular fate.

### Expression of early A/V lineage markers are recapitulated through EGFP lineage tracing within the cardiac crescent

Having identified a number of candidates that label atrial/ventricular cells at later stages of differentiation, we next investigated whether expression of any of these genes was associated with atrial/ventricular fate prior to formation of the chambers. The use of *Foxa2* linage-tracing information in our dataset allows us to identify the ventricular fated cells before they expressed markers such as *Irx4*, enabling profiling of ventricular progenitors at earlier stages than previous studies. We reasoned that profiling the *EGFP*+/- cells prior to tube or chamber formation might identify the genes and transcriptional programs that are turned on prior to acquisition of chamber-specific identity.

Towards that end, we performed clustering and gene expression analysis of cardiac regions of the CC (**Figure 5A**). We performed differential gene expression on a cluster-by-cluster basis and identified the top differentially expressed genes within each cluster, and used this information to identify several populations representing the *Pmp22+* LPM, *Isl1+/Tbx1+* aSHF, and the *Foxf1+/Aldh1a2+* pSHF **(Figure 5B/C).** We also identified a transcriptionally distinct *Hcn4+/Tbx5+* population representing the FHF. This population did not cluster separately in our merged analysis of all three timepoints (perhaps due to its concordance with differentiated cardiomyocytes at later stages), highlighting the utility in examining our data both across the merged data set and at each time point individually. *EGFP+* cells within the CC sample demonstrated that *Foxa2* lineage traced cells could be identified within particular progenitor populations at early stages of cardiogenesis within cells that have not yet taken on a bona fide atrial/ventricular transcriptional identity. In order to identify genes that were differentially regulated at this stage between atrial/ventricular fated cells, we performed differential gene expression analysis between *EGFP+ and EGFP-*cells within the heart field progenitors and LPM. We found differential expression of 101 genes which represent candidate markers for the earliest expressed genes involved in specification of the ventricular lineage **(Figure 5D).** GO enrichment analysis of the differentially expressed genes uncovered processes involved in lipoprotein biosynthesis and remodeling **(Figure 5E)**. A number of lipoproteins such as *Apoe* and *Apom* were differentially upregulated in the *EGFP+* cells. This was particularly interesting to us, as lipoproteins such as *Apobec2* were among the earliest upregulated genes along the ventricular trajectory construction previously (**Figure 4J**), and the role of metabolism remains poorly studied in early development. We further identified processes involved in cholesterol biosynthesis, suggesting that early metabolic differences may underlie the earliest segregation of atrial and ventricular cell types during development, potentially as early as the CC. Thus, through use of our genetic tools combined with scRNASeq technology we were able to identity a putative class of molecules involved in early adoption of ventricular identity. The spatiotemporal distribution of these lipoproteins at early stages of cardiac development has not been characterized extensively, thus future studies to understand the position and timing of lipoprotein expression within the cardiac crescent and developing heart tube are warranted.

**Figure 5.**
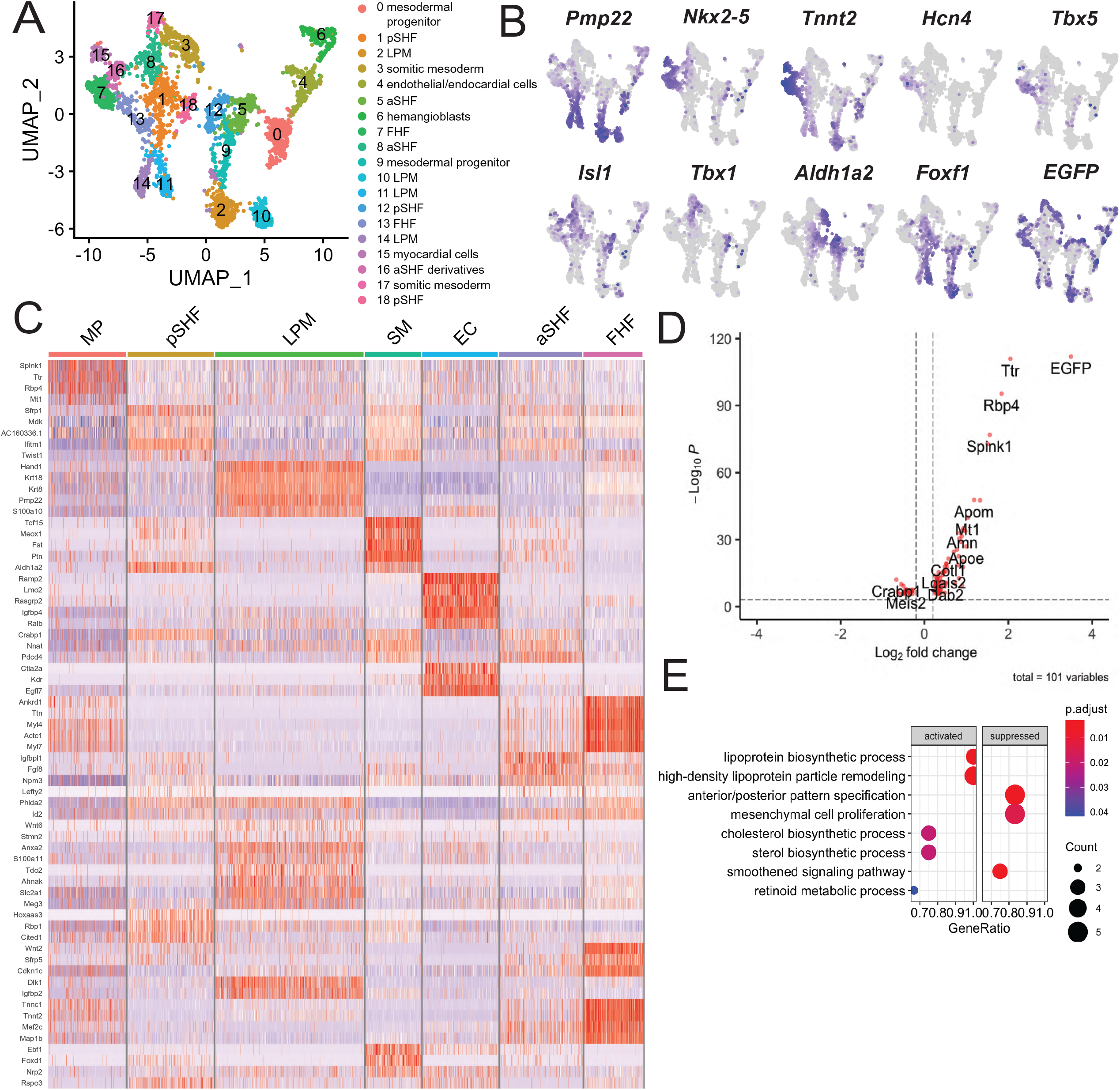
Characterization of Cardiac Crescent Populations and Identification of Transcriptomic Signature of Ventricular Fated Cell Types. **A)** UMAP clustering of cardiac cell types at the CC stage with cluster annotation. **B)** Feature plot for key markers of heart field progenitors. **C)** Heatmap demonstrating top 5 most differentially expressed genes for each cell type. Differentially expressed genes were identified using Wilcoxon rank-sum test with cutoff p-value <0.01. **D)** Volcano plot of differential expression across *EGFP+/-* cells within the CC. **E)** Selected upregulated/downregulated GO terms between *EGFP+/-* cells within the CC.

### Exogenous retinoic acid causes defects in head and heart development

While the sequential transcriptional mechanisms governing acquisition of chamber-specific fate are still not well understood, considerably more is known about the external cues that are necessary for proper development of the atrial and ventricular chambers. A number of previous studies have indicated that an anterior-posterior gradient of retinoic acid (RA) plays a role in directing differentiation of cardiac precursors towards an atrial fate (Bernheim and Meilhac, 2020; Perl and Waxman, 2019, 2020). Loss of *Raldh2* expression in the heart leads to absent synthesis of RA and loss of the atrial chambers (Niederreither et al., 2001). Exposure to exogenous RA *in utero* in turn is also associated with a number of heart and face defects, indicating that retinoids may act as teratogens during development of the heart (Piersma et al., 2017). Yet despite the recognition of the importance of RA signaling in these events, the direct downstream transcriptional effects induced by exogenous exposure to RA are incompletely understood in the context of chamber specification within the heart.

We and others have previously shown that manipulations in RA signaling during cardiac development leads to defects in FHF/SHF morphology in the CC as well as dysregulation of key components of the cardiac GRN, however the effect of RA exposure on the differentiation process at later stages of development were not completely explored (Bardot et al., 2017; Bertrand et al., 2011; Hochgreb et al., 2003; Lin et al., 2010; Rana et al., 2014; Stefanovic and Zaffran, 2017; Xavier-Neto et al., 1999). It is well established that the effects of RA are heavily dependent on its concentration, and that different concentrations can even have opposite effects (Perl and Waxman, 2019). To test the effect of RA gradients on early heart development we exposed mice to increasing concentrations of RA *in utero* at E7.25, and observed the effect of RA injection on embryonic development at E10.5. As expected, we find that exogenous RA causes patterning defects across the entire embryo in a dose-dependent manner (**Figure 6A**). Intraperitoneal injection of 65 mg/kg RA caused severe defects manifesting primarily in the head and pharyngeal structures, along with stunted formation of caudal structures. Exposure to 16.25 mg/kg RA caused milder defects, with grossly intact development of the tail and limb bud structures, with milder yet nonetheless hypomorphic effects on head and pharyngeal structures. Defects in heart development were also apparent, with dysregulation in chamber size within the developing heart tube. Low concentrations of RA (3.25 mg/kg) led to decreased size of the ventricle at E10.5 and mild craniofacial abnormalities. This indicated to us that this concentration was not overall detrimental to the embryo, and more suitable for understanding selective effects of exogenous RA signaling on formation of heart field progenitors and differentiation processes involved in morphogenesis of the heart chambers.

**Figure 6.**
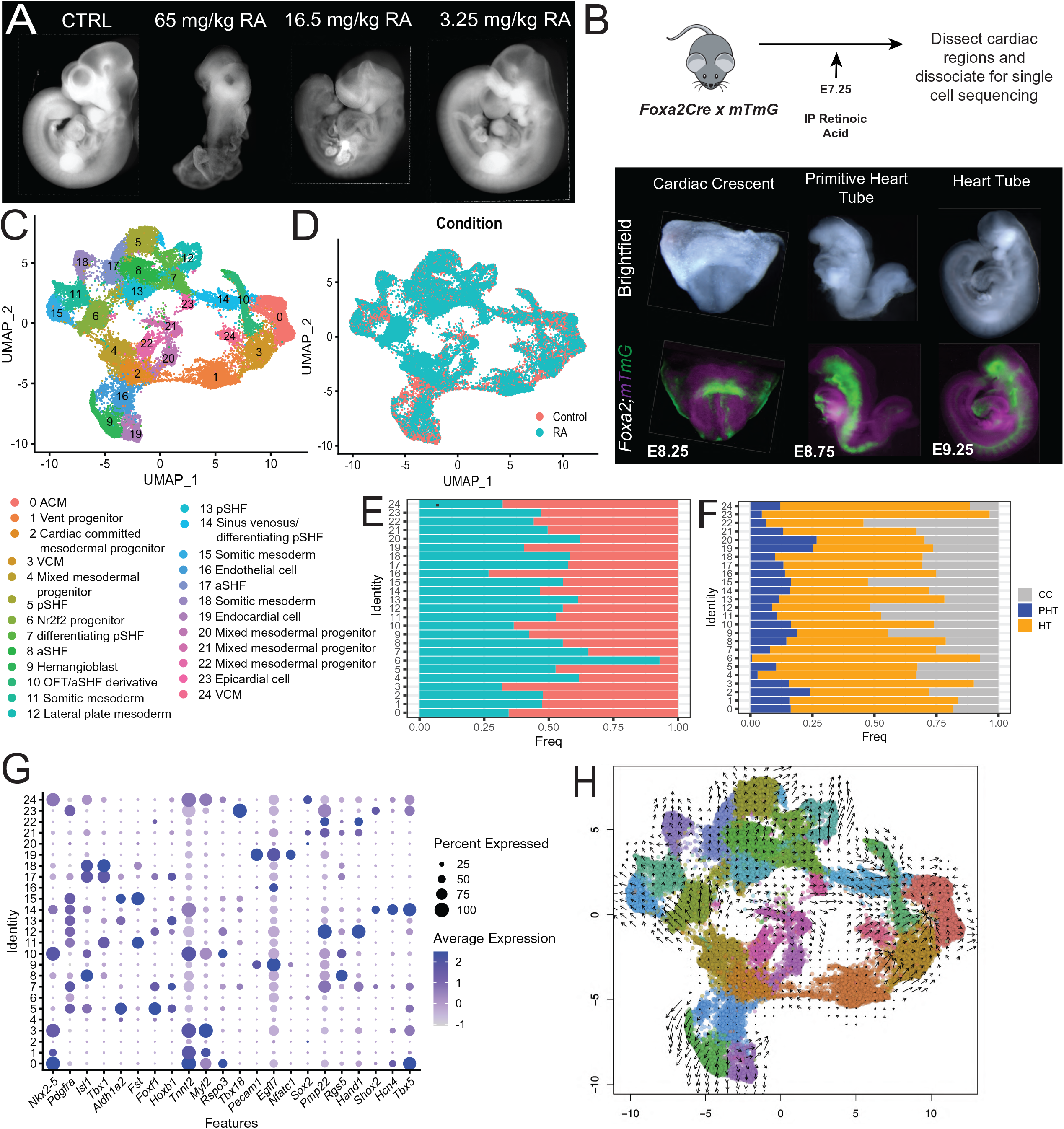
ScRNASeq following *in utero* Exposure to RA Demonstrates Defects in Ventricular Formation and Lineage Relationships Between aSHF and Ventricular Differentiation. **A)** Brightfield images of control and *in utero* exposed embryos demonstrating dose-dependent effects on heart and head development. B) *Top*: Experimental strategy for teratogenic mouse model of *in utero* RA exposure. *Bottom*: Representative images of RA-exposed embryos at stages sequenced. **C)** UMAP clustering of merged data set comprising control and RA-exposed embryos. **D)** Labeling of control (red) and RA (teal) cell types on UMAP embedding. **E)** Quantification of contribution to each cluster across control and RA samples. Frequency is calculated as total number of cells within cluster, relative to total number of cells within a treatment condition. **F)** Quantification of contribution to each cluster across embryonic time point samples. Frequency is calculated as total number of cells within cluster, relative to total number of cells within a sample. **G)** Bubble plot of selected markers across individual clusters. **H)** RNA velocity plot demonstrating directionality of differentiation across cell types.

In order to understand the downstream transcriptional effect of exogenous RA signaling in a cell-type specific manner, we injected pregnant dams (*Foxa2Cre;mTmG)* with 3.25 mg/kg RA intraperitoneally at E7.25. Embryos were collected at equivalent stages profiled previously from non-injected control embryos **(Figure 6B)**. We micro-dissected the cardiac regions of three stage-matched litter mates and sequenced >15,000 cells at each stage. We performed clustering and differential expression to annotate the different cell types present at each stage using similar methods described above, followed by sub-setting the cardiac regions of each sample using equivalent criteria as those applied to non-injected hearts.

We merged samples from CC, PHT and HT stage embryos across control and RA-injected embryos and performed dimensionality reduction and clustering as done previously (**Figure 6C/D**). We observed that cells from control and RA-injected embryos contributed to all clusters and that the UMAP projections remained similar to the control samples of the earlier analysis. This indicated to us that gross differences in chamber formation and cell identity were not induced by low concentrations of RA, consistent with the mild morphological changes observed (**Figure 6A**). We performed differential gene expression in order to annotate cluster identity, and quantified the frequency of sample contribution to each cell type for both WT and RA conditions (**Figure 6G**). Similar to our previous data from control embryos, we saw differential contribution of cells from early stage (CC) and late stage samples (HT) to particular populations in a manner that recapitulated the growth and differentiation of these processes (**Figure 6F**).

### Exogenous retinoic acid causes defects in contribution to ventricular cells, but not heart field progenitors

Given the defects in FHF/SHF morphology and size in previously published experiments, we first tested if exogenous RA affected the contribution of cell types to clusters of aSHF or pSHF identity. We observed no differences in the relative proportion of cells from control or RA-injected embryos on aSHF and pSHF identity, nor with the LPM population (**Figure 6E**). This indicated to us that at this concentration, exogenous RA did not influence the relative size of aSHF or pSHF populations. The overall frequency of cells that contribute to the ventricular cardiomyocytes (cluster 3) however was decreased in RA injected embryos relative to the size of all other cells in the samples. This was in line with our observation that exogenous RA caused defects in the size of the ventricular chambers macroscopically. While the majority of the UMAP projections appeared similar to those in the control embryos, there was notable dysconnectivity between the aSHF cells (clusters 8,17) and the differentiating aSHF derivatives (cluster 10) which cluster adjacent to ventricular cells **(Figure 6C).** This led us to ask whether defects in differentiation of aSHF cell types towards a cardiomyocyte lineage was affected by exogenous RA, even if the relative size of the aSHF population was not impacted. We visualized RNA velocity trajectories across the various populations from both control and RA-injected embryos **(Figure 6H)**. While pSHF populations appeared to differentiate rapidly into atrial cells (cluster 0) as observed in control embryos, the differentiation of aSHF cells appeared to deviate away from ventricular cells and converged toward a separate cluster in proximity to the somitic mesoderm (cluster 6). This cluster was also overwhelmingly overrepresented in RA samples, indicating that exogenous RA had caused defects in the normal lineage trajectory of aSHF progenitors, resulting in a shunt of these cells toward a somitic precursor cell type (**Figure 6E)**. Cluster 10 (representing differentiating aSHF cell types) also had decreased contribution of cells from RA-exposed samples. Collectively, our data indicates that *in utero* exposure to RA had an inhibitory effect on ventricular size by disrupting the differentiation of aSHF cells towards their usual ventricular progeny. Given the multipotent nature of the aSHF, it is perhaps not surprising that defects in ventricular differentiation would lead to increased contribution to pharyngeal lineages. However, it is not as of yet clear if this is caused by overexpression of genes involved in somitic mesoderm differentiation as a consequence of RA injection, or if this represents a compensatory mechanism to account for failure to differentiate along a ventricular lineage.

### RA causes dysregulation in processes guiding ventricular specification of progenitor populations

We next wished to more comprehensively understand which changes in the transcriptome of RA-exposed cells drove the failure to differentiate progenitors along a ventricular lineage, and to what extent changes in the transcriptome were cell-type dependent. Differential expression analysis within ventricular cells across RA and control cell types identified 4,128 differentially expressed genes. RA injection caused decreases in processes involved in striated muscle development and ventricular muscle morphogenesis, but also caused defects in metabolic and energetic components including components of the respiratory electron transport chain (**Figure 7A**). Other genes that were downregulated in the RA condition included general sarcomeric genes and cardiac transcription factors such as *Nkx2-5* and the *Foxa2* lineage tracing marker *EGFP,* as well as a number of bona fide ventricular markers such as *Myl2 and Ckb* **(Figure 7A).** These processes indicated that not only were ventricular cells present at lower frequency, they also appeared comparatively less healthy. Interestingly, ventricular cells from RA-injected embryos expressed higher levels of a number of atrial markers and insulin-like growth factor signaling, underlying the improper specification of atrial/ventricular identity.

**Figure 7.**
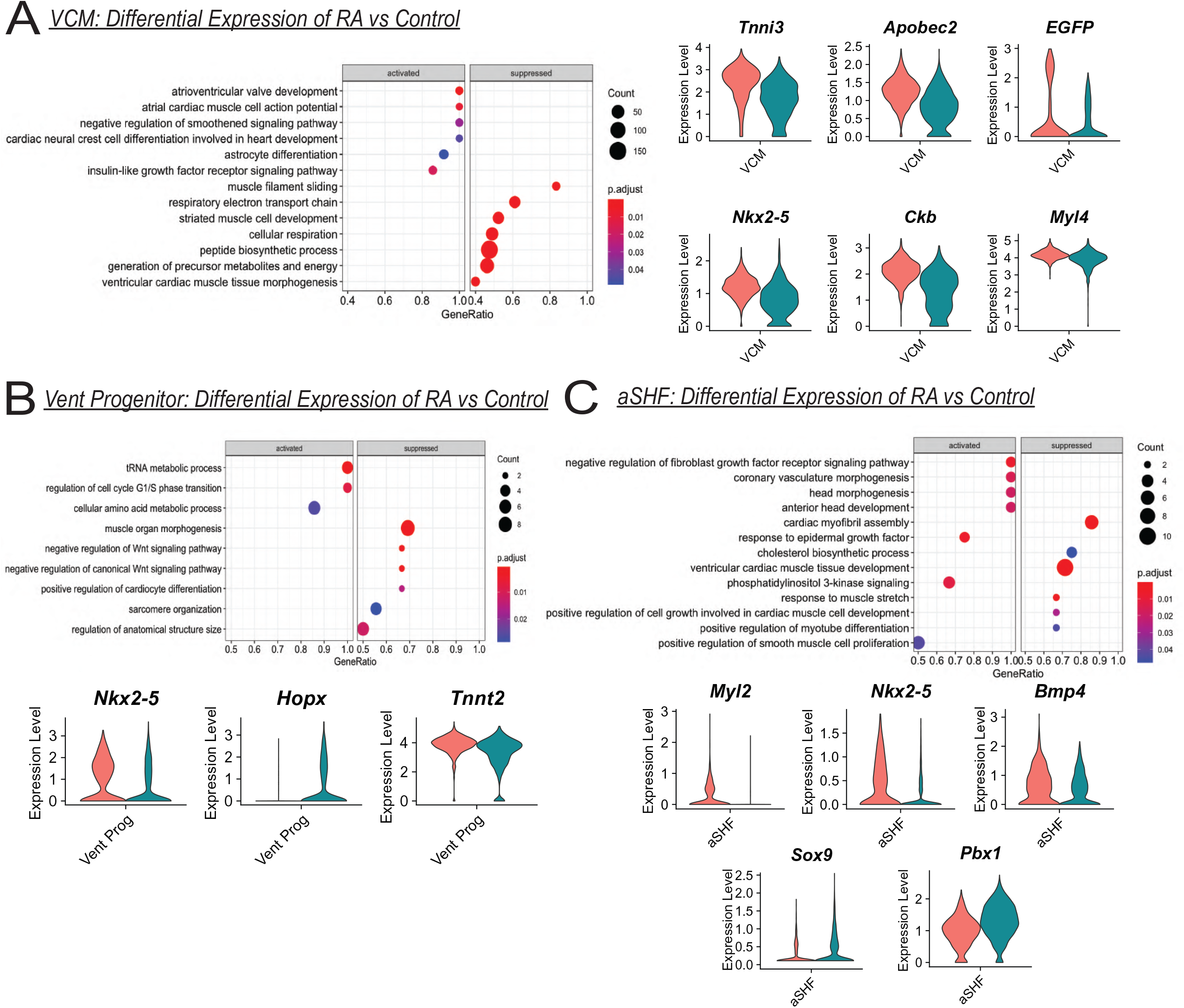
Differential Expression of Ventricular Cells and Associated Progenitors Shows Dysregulation in Processes Related to Ventricular Development and Differentiation. **A)** *Left:* Selected up/downregulated GO terms within RA exposed ventricular cardiomyocytes *Right:* Selected differentially expressed candidates between control (red) and RA (teal) cells within ventricular cluster. B) *Top:* Selected up/downregulated GO terms within RA-exposed early myocardial progenitors; *Bottom:* Selected differentially expressed candidates between control (red) and RA (teal) cells within early myocardial progenitors. C) *Top:* Selected up/downregulated GO terms within RA-exposed aSHF cluster; *Bottom:* Selected differentially expressed candidates between control (red) and RA (teal) cells within aSHF cluster.

To assess whether similar processes were affected in ventricular progenitors we performed differential gene expression and gene set enrichment analysis, which uncovered increased expression of processes involved in biosynthetic machinery such as “tRNA metabolism” and “amino acid synthesis” (**Figure 7B**). This was coupled with decreased sarcomere organization and differentiation toward the cardiomyocyte lineage accompanied by dysregulation in Wnt signaling, suggesting that these cells were arrested during differentiation towards a cardiac lineage. The progenitors in RA-injected embryos demonstrated decreased *Nkx2-5* and *Tnnt2* expression but interestingly had increased expression of *Hopx,* perhaps indicative of an arrest of differentiation at the cardiomyoblast stage that could not be overcome to form functional cardiomyocytes (Jain et al., 2015).

Lastly, we set out to understand processes that were differentially regulated within the aSHF that might have caused these defects. We performed gene set enrichment on differentially expressed genes between RA-injected and control embryos and found that exogenous RA caused a downregulation in processes involved in “myotube development”, “ventricular cardiac muscle development” and “cardiac myofibril assembly”, as well as decrease in “cholesterol synthesis”. These were accompanied by an increase in processes involved in head morphogenesis and head development consistent with the subtle head deformities that had been observed prior. We also uncovered increased negative regulation of FGF signaling, which is known to be required for differentiation of aSHF cells towards the cardiomyocyte lineage (De Zoysa et al., 2020; Pradhan et al., 2017). Consistent with this, aSHF cells from RA-exposed embryos had decreased expression of cardiomyocyte markers such as *Myl2* and *Nkx2-5* with downregulation of *Bmp4* which plays a key role in differentiation of the aSHF progenitors. These changes were accompanied with increases in *Pbx1* and *Sox9,* both genes involved in development of the pharyngeal lineages (Cheung and Briscoe, 2003; Welsh et al., 2018). These data suggest that exogenous RA caused defects in different signaling pathways that subsequently caused a block in differentiation of aSHF towards a ventricular lineage and may have caused a shunt towards other lineages within its multipotent capacity, such as the somitic mesoderm.

Taken together, exogenous RA appeared to cause defects in ventricular differentiation that affected both expression of key pathways required for differentiation of the aSHF such as FGF signaling, as well as having profound effects on expansion and differentiation of cardiac progenitors. Future work will need to validate these findings in more depth, paying particular attention to whether these differences arise because of metabolic requirements for ventricular cells as they differentiate, or if they represent early processes that aid in specification and commitment toward the ventricular lineage. Further follow up analysis of this data as well as functional follow up of these processes will be needed to fully understand the role of metabolism in differentiation and lineage specification and commitment.

## Discussion

The complex lineage relationships between cardiac progenitors and their differentiated progeny during the earliest steps of commitment toward a cardiac lineage and formation of the heart chambers remains of active interest, not least due to the notion that many congenital heart defects are thought to arise at this critical time of development (Francou and Kelly, 2016; Kloesel et al., 2016). By conducting scRNASeq of multiple timepoints from the beginning of cardiac specification to early chamber morphogenesis, we have generated a map of the dynamic transcriptional landscape of the heart.

One of our primary interests was to elucidate the molecular mechanisms of early atrial and ventricular lineage specification. Based on previous work by us and others, we utilized *Foxa2* lineage tracing to capture the ventricular cells throughout this temporal analysis. This is particularly relevant prior to chamber formation, and prior to expression of the canonical chamber-specific markers. Our analysis identified multiple known cardiac progenitor populations including the FHF/aSHF/pSHF and LPM progenitors and identified their molecular signature, as well as the differences between them. We also identified a separate population of *Pmp22+/Nkx2-5+* mesodermal population that clustered separately from the LPM together with endothelial/endocardial cell types and appeared to give rise to a mixed population of atrial and ventricular cardiomyocytes. Along these lines, we identified known and new candidate genes expressed in these distinct populations, and established a system to interrogate the changes that occur over time of differentiation toward more differentiated cell types. These data help to build on existing work defining the heterogeneity of heart field progenitors (Ivanovitch et al., 2021; de Soysa et al., 2019; Tyser et al., 2021) by relating them transcriptionally to their more differentiated progeny.

Towards that end, our data captured discrete clusters of SHF progenitors at multiple stages of differentiation, including their “stable” replicative state as well as intermediary states during differentiation towards a cardiomyocyte/pharyngeal lineage. Use of RNA velocity tools helped us to identify rapidly differentiating cells and the lineage relationships between progenitors and their progeny in an unbiased manner. While previous studies have profiled the heterogeneity of heart field progenitors at similar stages (Ivanovitch et al., 2021; de Soysa et al., 2019; Tyser et al., 2021), few others have conclusively identified these intermediary differentiating states and used them to interrogate differences in signaling pathways between cardiac progenitors during differentiation in a comprehensive manner. This analysis identified several previously described concepts -- for instance the upregulation of *Fgf8* and *Bmp4* in differentiation of the aSHF toward the OFT/RV -- but also demonstrated differences in canonical/non-canonical Wnt signaling across aSHF/pSHF differentiation events. We also observed that various different components of the cardiac GRN such as Tbx5/Tbx20 and other processes involved in early differentiation of heart progenitors such as Robo/Slit signaling were differentially expressed across these two differentiation streams.

We show that while FHF cells were identified as a discrete population within the CC stage, these cells became integrated together with other differentiated cardiomyocytes once merged with samples from later stages. These data add to a growing understanding of the FHF as a spatially distinct region within the cardiac crescent whose molecular signature resembles that of early differentiating cardiomyocytes and is altogether different from its SHF counterpart.

We were surprised to find that the differentiation streams did not appear to separate entirely on the basis of eventual atrial/ventricular fate, as we identified differentiating *EGFP+* ventricular fated cells from the LPM clustered together with differentiating cells from the pSHF towards the SV and atrial lineages. This suggests the possibility that utilization of similar pathways and regulators of cardiac fate such as Tbx5/Tbx20 and the Gata factors might in fact be shared across different progenitors (such as the FHF and the pSHF) to drive differentiation towards different chamber fates, though further work would be needed to confirm this hypothesis. If true, the molecular basis of how similar programs could be used to instruct differentiation toward both atrial and ventricular cells would be of significant interest – these context-specific outcomes could be driven by epigenetic differences or spatiotemporal segregation of progenitors that alter how cells respond to shared cues from their environment.

We identified several distinct populations of atrial and ventricular cells within our data, and performed differential expression analysis to identify putative candidates of atrial/ventricular fate and identity. We also observed that atrial and ventricular cells appeared to arise from multiple separate progenitor cell types including our newly identified *Pmp22+/Nkx2-5+* population, suggesting that convergent differentiation processes may drive the formation and growth of the chambers. Further work will be needed to understand if segregation of these lineages from different progenitors occurs through the same mechanisms as those governing differentiation of cardiomyocytes from aSHF/pSHF/LPM progenitors. By combining *Foxa2* lineage tracing with single cell sequencing technology, we were also able to identify genes involved in the earliest stages of differentiation toward a ventricular lineage at the cardiac crescent stage. This identified several candidate genes, including a number of lipoproteins that have not previously been described to be involved in chamber development.

Lastly, we utilized a teratogenic model of *in utero* exposure to RA to study how RA impacts the signaling landscape and resulting cellular composition of the early heart, particularly the mechanisms relating to atrial/ventricular differentiation. We find that exogenous RA leads to the formation of heart and head defects in a dose-dependent manner. At low doses, RA caused formation of hypomorphic ventricular chambers. Ventricular cells from RA-exposed embryos had aberrant expression of atrial-like processes and expressed processes indicative of reduced energetic fitness due to defects in mitochondrial ATP generation and sarcomere assembly. Exogenous RA also caused impaired differentiation of aSHF cells towards a cardiomyocyte lineage, leading to increased expression of neural crest markers involved in head/pharyngeal development that accompanied decreased expression of cardiomyocyte differentiation markers within the aSHF. Future results will examine whether RA exposure causes ventricular defects at later stages in a right/left specific manner due to the involvement of the aSHF in this process, as well as more comprehensively characterize facial defects arising from increased specification of somitic mesodermal cell types. Further studies into these processes will help us not only understand the pathological mechanisms of this teratogenic model, but will further contribute to our understanding of targets downstream of RA signaling in a cell type-specific manner and with a focus on specification of atrial and ventricular cells during normal development.

## Materials and Methods

### Mice

Foxa2Cre mice (C57BL/6J) mice were kindly shared with us by Dr. Heiko Lickert (Horn et al 2021). Rosa26-mTMG mice were obtained from the Jackson Laboratory. Studies were performed on mixed backgrounds depending on strain availability. All murine embryo scRNASeq studies were performed on Foxa2Cre;mTmg embryos derived from crossing Foxa2Cre males with Rosa26-mTmG females. For timed matings, the day of plug identification was considered to be E0.5. All animals were housed in the Center for Comparative Medicine and Surgery (CCMS) facilities at Icahn School of Medicine and all experiments were conducted in accordance with the guidelines and approval of the institutional Animal Care and Use Committee at Icahn School of Medicine at Mount Sinai. For studies on the effect of retinoic acid, pregnant mTmG females previously crossed with Foxa2Cre males were injected intraperitoneally at E7.25 with 65 mg/kg, 16.25 mg/kg, or 3.25 mg/kg of retinoic acid dissolved in corn oil. Single cell analysis on RA-exposed embryos was performed following injection of 3.25 mg/kg retinoic acid.

### Dissection and isolation of murine cardiac tissues

Foxa2Cre;mTmg embryos were dissected at E8.25, E8.75 and E9.25 and identified . The cardiac region was sub-dissected at each stage (see Supplementary Figure 1), including the surrounding endoderm, head folds, and pharyngeal structures. This was done in order to ensure complete dissection of the cardiac structures and also to include transcriptional information within key tissues that co-develop with the cardiac structures at each stage. Embryos were dissected in cold PBS with 0.1% BSA and kept on ice until dissociation. Three stage matched litter-mates were incubated separately in 200 ul of 0.25% Trypsin-EDTA in a 37C water bath for 5 minutes or until the tissue dissociated completely, vortexing once halfway. Trypsin solution was quenched with 1ml DMEM with 10% FBS, centrifuged at 300g for 5 minutes, and resuspended in 50 ul PBS with 0.1% BSA for single-cell RNA sequencing preparation.

### Single cell cDNA library preparation and scRNA sequencing

Individual samples for each time point were hashed using the Chromium platform in order to determine that downstream analysis was not affected by biological variability between embryos. After hashing, libraries were generated using the Chromium platform with the 3’ gene expression (3’ GEX) V3 kit, using an input of ∼10,000 cells. Gel-Bead in Emulsions (GEMs) were generated on the sample chip in the Chromium controller. Barcoded cDNA was extracted from the GEMs by Post-GEM RT-cleanup and amplified for 12 cycles. Amplified cDNA was then fragmented and subjected to end-repair, poly A-tailing, adapter ligation, and 10X-specific sample indexing following the manufacturer’s protocol. Libraries were quantified using Bioanalyzer and QuBit analysis. Libraries were sequenced in paired end mode on a NovaSeq instrument targeting a depth of 50,000 reads per cell.

### Processing of sequencing reads

Sequencing data was aligned and quantified using the Cell Ranger Single-Cell Software Suite (version 3.0, 10x Genomics) against the provided GRCm38 (mm10) mouse reference genome. Lineage tracing information was determined by further alignment to a custom reference for EGFP using provided maps of constructs used in creation of *mTmG* mice from Jackson Laboratory. Complimentary Tomato expression information could not be obtained due to increased distance from the polyA tail compared to EGFP.

### Clustering and differential expression analysis

Downstream differential expression and clustering analysis was performed using the Seurat V.4.0 package, as described in the tutorials (http://satijalab.org/seruat/). *cellRanger* matrices were imported for each sample, and distributions of nCountRNA, nFeatures and % mitochondrial gene expression were examined for each sample in order to filter out doublets or cells of low quality. Cells with greater than 20% of genes coming from mitochondrial genes were selected against, as well as those with fewer than 200 genes.

We then normalized the resulting subset Seurat objects using the scTransform workflow and further scaled and normalized the RNA assay in order to perform downstream differential expression analysis. We performed principal component analysis using the highly variable genes for each sample. The most significant principal components (20-30 depending on sample) were used for graph-based unsupervised clustering (FindClusters and FindNeighbor functions). Uniform manifold approximation and projection (UMAP) was performed using a standard resolution parameter of 0.5 and iteratively modified after performing marker gene expression and examining expression of key markers.

In order to generate a combined Seurat object encompassing all three timepoints, we visualized expression of cardiac and mesodermal genes (*Pdgfra, Nkx2-5*), endothelial/endocardial (*Pecam-1, Nfatc1*), endodermal (*Epcam, Sox17*), and ectoderm (*Sox2, Pou3f1*). We subset clusters with positive expression of mesodermal/cardiac markers, and more completely annotated clusters that did not express these markers but clustered closely through GO/KEGG analysis and differential expression analysis. These represented related lineages such as the neural crest or early/related mesodermal progenitors. These were subset together with the cardiac lineages for each stage. Seurat objects for the CC, PHT and HT stage were labeled with their original sample ID and merged together using the scTransform based integration workflow. For analyses involving comparison of Normal and RA-injected embryos, the above steps were repeated by combining objects from both conditions either for corresponding stages or across all samples. Downstream clustering and dimensionality reduction was performed using similar methods described above. Cell Cycle quantification and annotation was performed on merged samples using the CellCycleScoring vignette available from the Satija lab. Quantification of time point, condition, or sample contribution to clusters was determined by normalizing the frequency of a given metadata classification to the total cell number from each sample in order to control for cell number differences across samples.

In most cases, differential expression was determined using the FindMarkers function on the “RNA” assay, utilizing a negative binomial model with parameters for min.pct = 0.2 and a p-value cut off of 0.01 either by comparing a single cluster to all others for annotation, or in pairwise fashion for clusters of interest. Differential expression of Foxa2-lineage traced positive and negative cells was done by classifying each cell with a binary variable based on expression of EGFP > 1.0 and then performing differential expression on subsets of cells across the EGFP +/- metadata information. A similar strategy was employed to query differentially expressed genes due to RA injection on a cluster-by-cluster basis across Normal and RA condition metadata information.

### Gene set enrichment and GO/KEGG analysis

Differential gene expression was performed in Seurat through the FindMarkers function in Seurat using a negative binomial test using a p-value cutoff of < 0.05 as described previously. Gene set enrichment for GO terms and KEGG pathways were performed using the ClusterProfiler tool (https://guangchuangyu.github.io/software/clusterProfiler/) once again using a cutoff p-value of <0.05.

### Atrioventricular lineage trajectory visualization

Atrial and ventricular trajectories were visualized using Monocle3 (https://cole-trapnell-lab.github.io/monocle3/) following importation of various parameters from Seurat (UMAP embedding, RNA counts, etc). Trajectories were learned utilizing the learn_graph function in Monocle, with the following parameters: ncenter= 350; rann.k = 25; minimal_branch_len = 15). Cells were ordered utilizing marker expression for progenitors and differentiated cell types, as well as quantification of cell type contribution from early vs late sample (i.e. cardiac crescent stage vs heart tube).

### RNA velocity analysis

RNA velocity was calculated using VelocytoR according to the developer’s manual. Briefly, .loom files containing exon and intron information of all samples were created using the Velocyto python package *velocyto run10x* command and merged using *loompy.combine* in the loompy package (http://loompy.org). All subsequent analysis was performed in R. To incorporate splicing information into previously created Seurat objects, .loom files were subset and filtered accordingly and added as separate assays using the Seurat *CreateAssayObject()* command. RNA velocity was then calculated using the *RunVelocity()* command in the SeuratWrappers package and visualized on the previously computed UMAP embedding using *show.velocity.on.embedding.cor()* in the velocyto R wrapper package velocyto.R.

### Data availability

Single cell RNA sequencing data (count matrices/metadata and raw sequencing data) will be deposited in the NCBI Gene Expression Omnibus upon publishing of this work.

## Supplementary Figure Legends

**Supplementary Figure 1.**
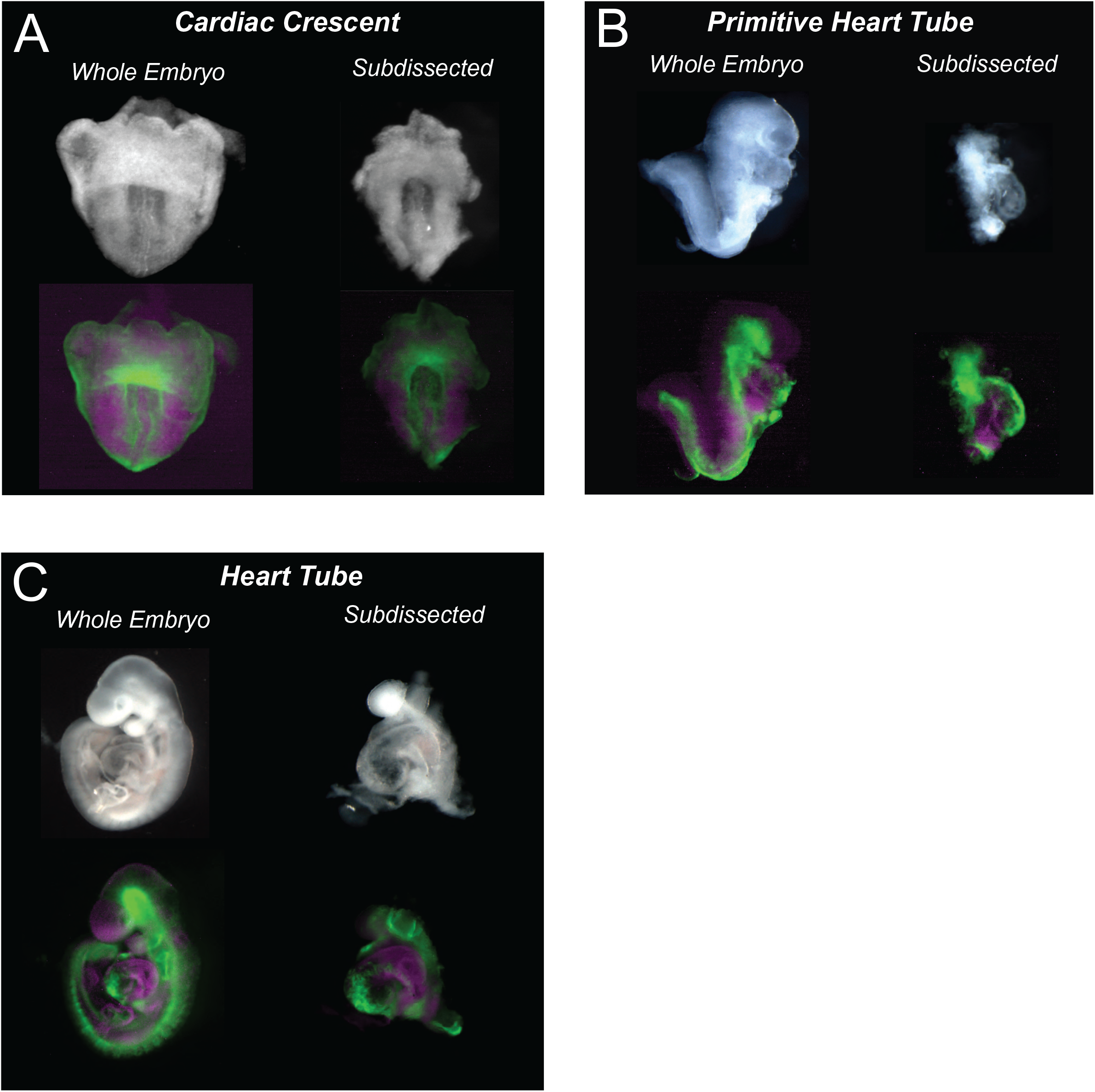
Representative Images of Embryonic Stages Used for scSeq and Dissection Examples. **A)** Cardiac Crescent stage (E8.25) embryo showing brightfield (top) and Foxa2-mTmG fluorescence (bottom). Left = whole embryo; right = example dissection of cardiac regions **B)** Primitive Heart Tube (E8.75) stage embryo showing brightfield (top) and Foxa2-mTmG fluorescence (bottom). Left = whole embryo; right = example dissection of cardiac regions **C)** heart Tube stage (E9.25) embryo showing brightfield (top) and Foxa2-mTmG fluorescence (bottom). Left = whole embryo; right = example dissection of cardiac regions.

**Supplementary Figure 2.**
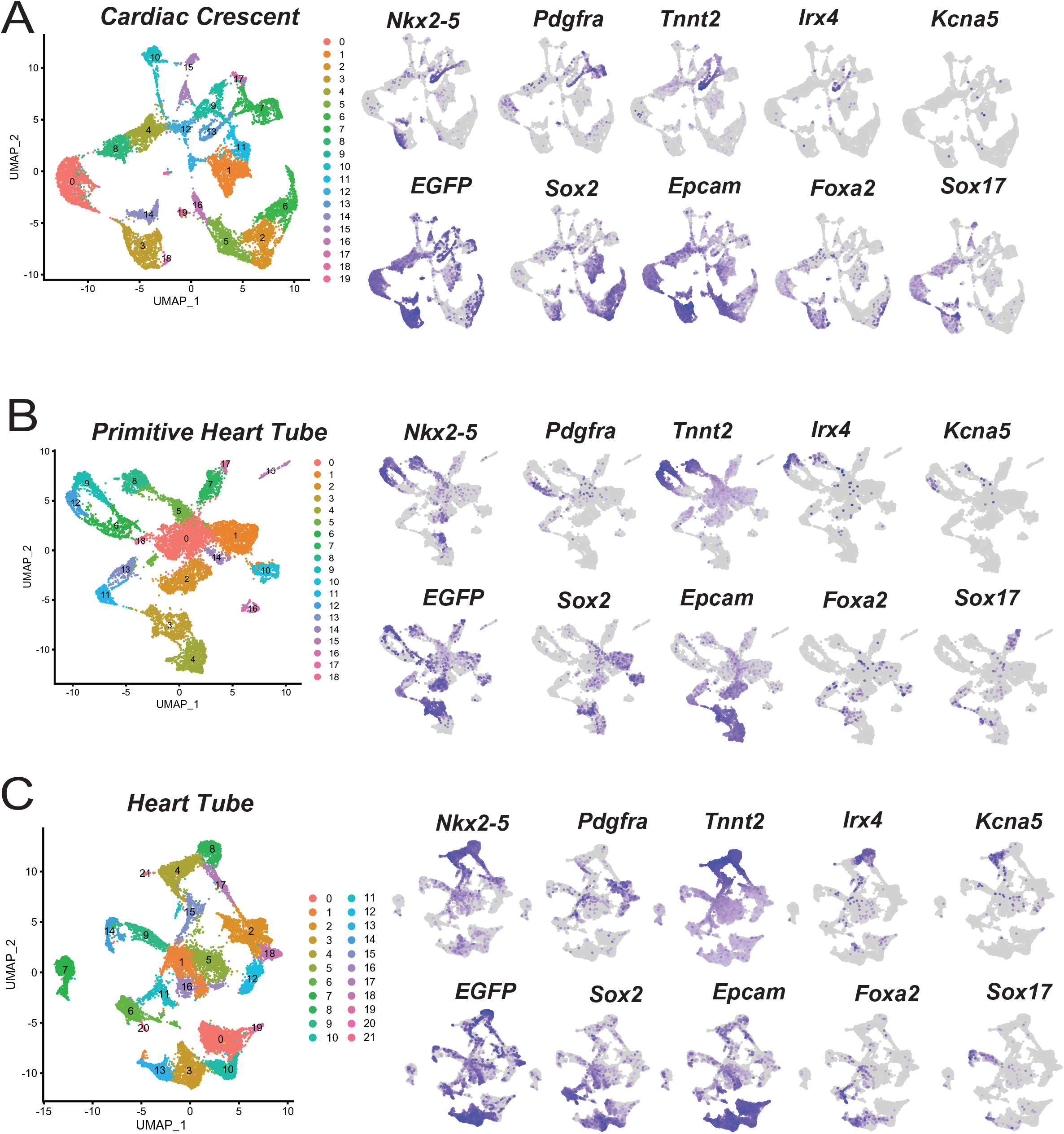
UMAP Clustering and Selection of Key Markers for Populations of Interest. **A)** UMAP clustering and feature plots for markers of mesodermal, endodermal, and ectodermal cell types, as well as atrial/ventricular cells at CC stage. **B)** UMAP clustering and feature plots for markers of mesodermal, endodermal, and ectodermal cell types, as well as atrial/ventricular cells at PHT stage. **C)** UMAP clustering and feature plots for markers of mesodermal, endodermal, and ectodermal cell types, as well as atrial/ventricular cells at HT stage.

**Supplementary Figure 3.**
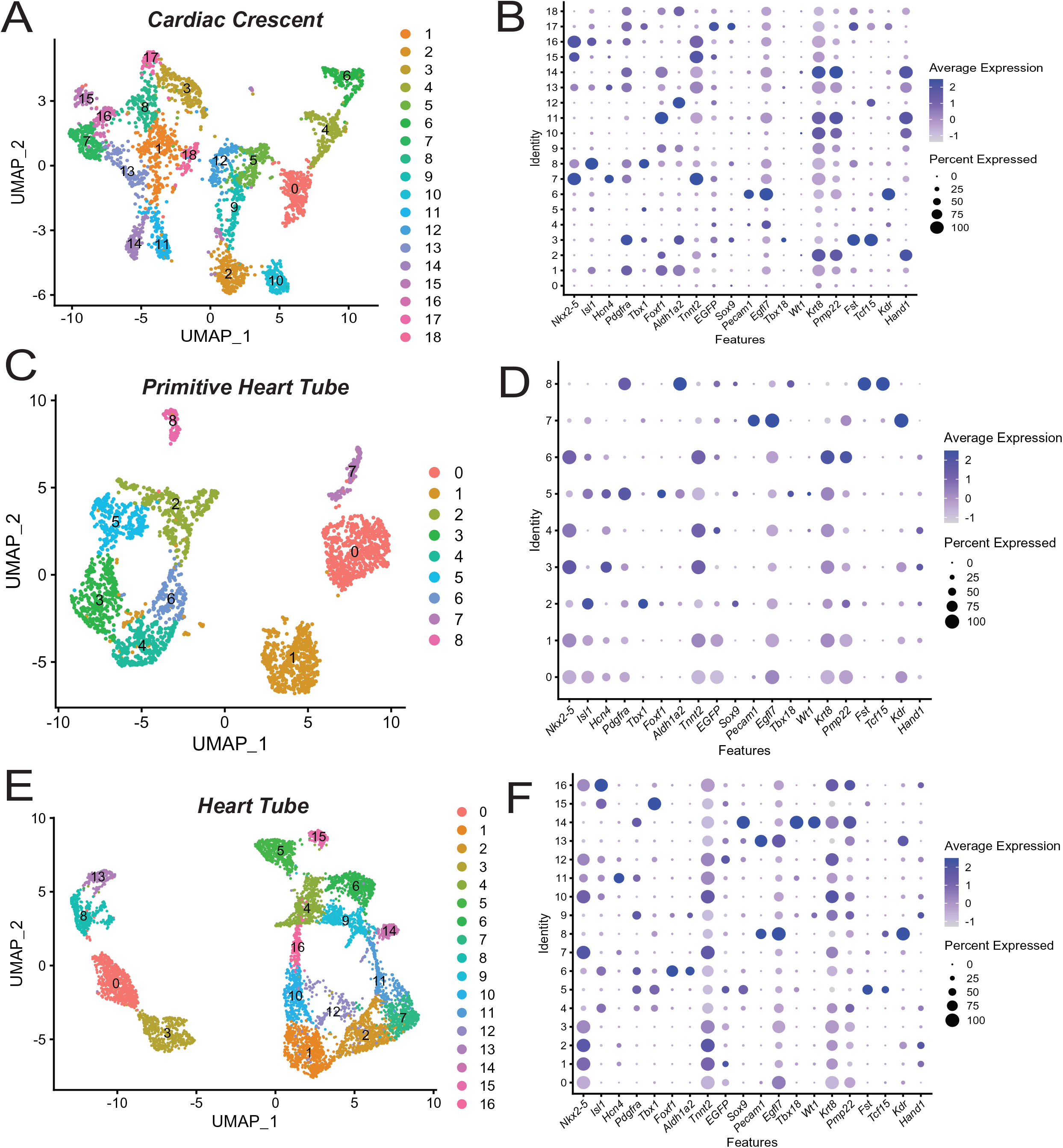
UMAP Clustering and Selection of Key Markers for Cardiac Subpopulations at Individual Stages. **A)** UMAP clustering of subclustered cardiac cells at CC stage. **B)** Dot plot of selected genes of interest at PHT stage. **C)** UMAP clustering of subclustered cardiac cells at PHT stage. **D)** Dot plot of selected genes of interest at PHT stage. **E)** UMAP clustering of subclustered cardiac cells at HT stage. **F)** Dot plot of selected genes of interest at HT stage.

**Supplementary Figure 4.**
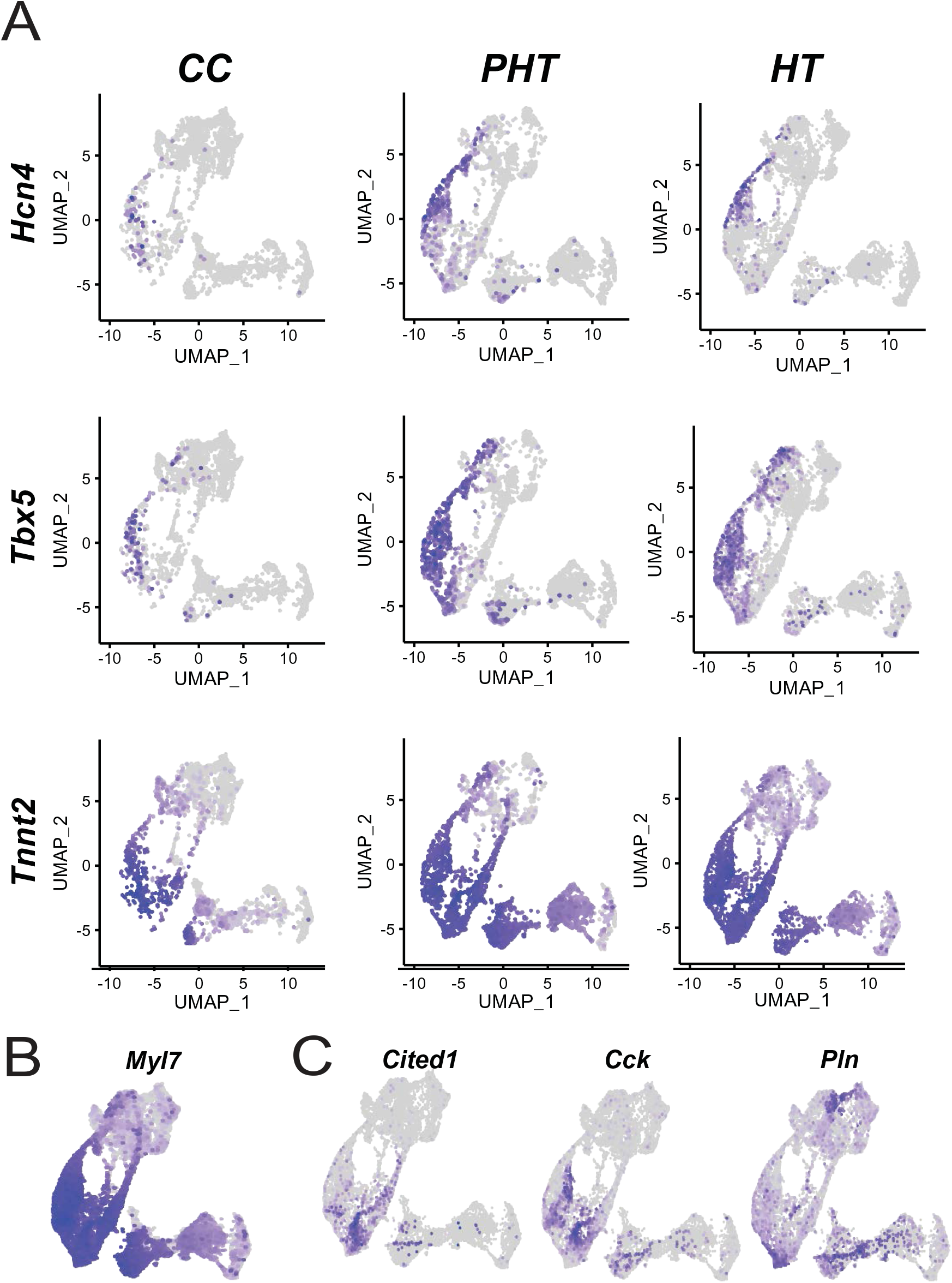
Expression of Key Markers of Interest Across Multiple Stages of Development. **A)** Feature plots for *Hcn4, Tbx5,* and *Tnnt2* split separately across individual samples. **B)** Feature plot for Myl7 expression. **C)** Feature plots for expression of right (*Cited1/Cck)* and left (*Hand1/Pln)* ventricular identity.

**Supplementary Figure 5.**
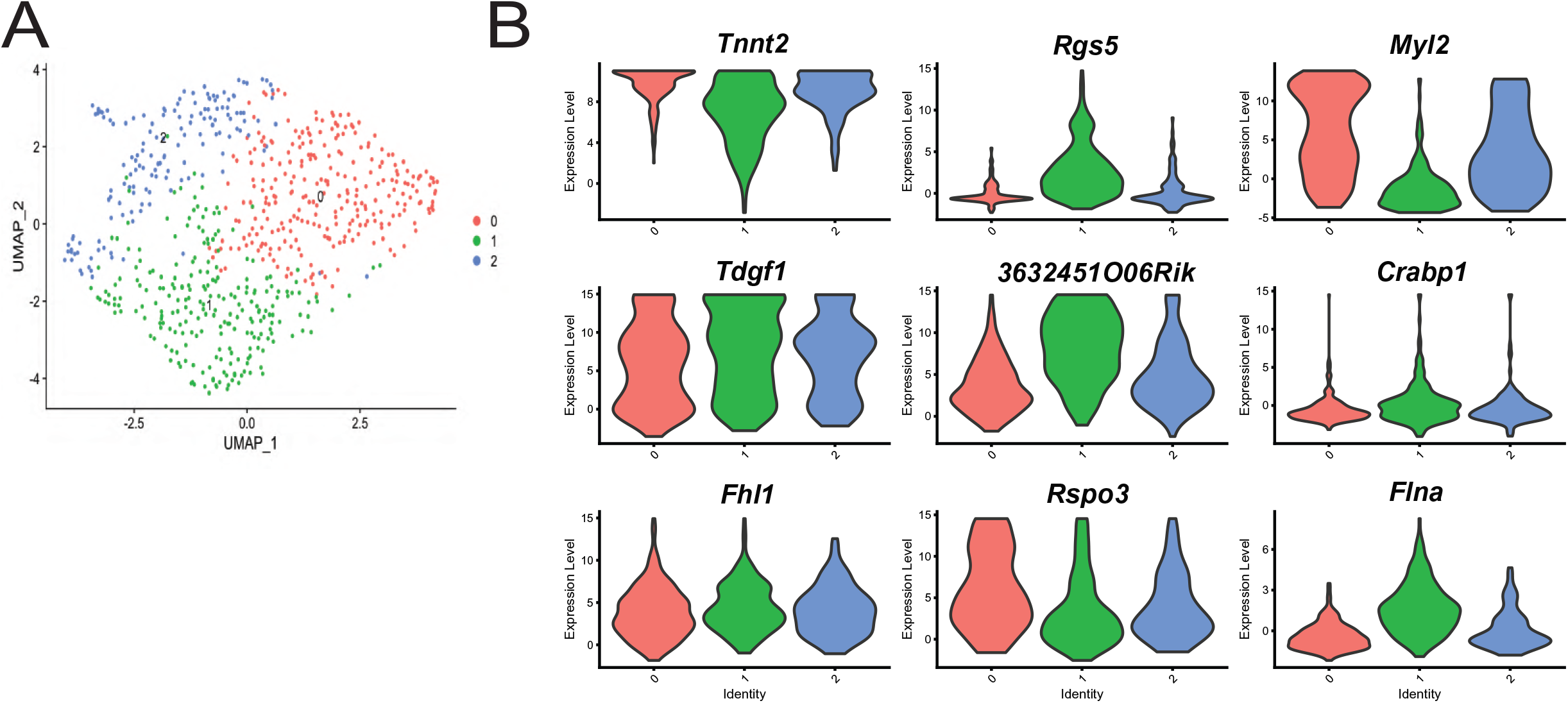
Expression of OFT/Ventricular Markers Following Subclustering of Cluster 5. **A)** UMAP clustering of Cluster 5 alone following re-clustering. **B)** Violin plots for markers of ventricular and OFT lineages across individual clusters.

## References

Bardot, E., Calderon, D., Santoriello, F., Han, S., Cheung, K., Jadhav, B., Burtscher, I., Artap, S., Jain, R., Epstein, J., et al. (2017). Foxa2 identifies a cardiac progenitor population with ventricular differentiation potential. Nat Commun 8, 14428.

Bardot, E.S., Jadhav, B., Wickramasinghe, N., Rezza, A., Rendl, M., Sharp, A.J., and Dubois, N.C. (2020). Notch Signaling Commits Mesoderm to the Cardiac Lineage. BioRxiv 2020.02.20.958348.

Bernheim, S., and Meilhac, S.M. (2020). Mesoderm patterning by a dynamic gradient of retinoic acid signalling. Philos Trans R Soc Lond B Biol Sci 375, 20190556.

Bertrand, N., Roux, M., Ryckebüsch, L., Niederreither, K., Dollé, P., Moon, A., Capecchi, M., and Zaffran, S. (2011). Hox genes define distinct progenitor sub-domains within the second heart field. Developmental Biology 353, 266–274.

Chabab, S., Lescroart, F., Rulands, S., Mathiah, N., Simons, B.D., and Blanpain, C. (2016). Uncovering the Number and Clonal Dynamics of Mesp1 Progenitors during Heart Morphogenesis. Cell Rep 14, 1–10.

Cheung, M., and Briscoe, J. (2003). Neural crest development is regulated by the transcription factor Sox9. Development 130, 5681–5693.

Chiapparo, G., Lin, X., Lescroart, F., Chabab, S., Paulissen, C., Pitisci, L., Bondue, A., and Blanpain, C. (2016). Mesp1 controls the speed, polarity, and directionality of cardiovascular progenitor migration. J Cell Biol 213, 463–477.

De Zoysa, P., Liu, J., Toubat, O., Choi, J., Moon, A., Gill, P.S., Duarte, A., Sucov, H.M., and Kumar, S.R. (2020). Delta-like ligand 4-mediated Notch signaling controls proliferation of second heart field progenitor cells by regulating Fgf8 expression. Development 147, dev185249.

DeLaughter, D.M., Bick, A.G., Wakimoto, H., McKean, D., Gorham, J.M., Kathiriya, I.S., Hinson, J.T., Homsy, J., Gray, J., Pu, W., et al. (2016). Single-Cell Resolution of Temporal Gene Expression during Heart Development. Dev Cell 39, 480–490.

Devalla, H.D., Schwach, V., Ford, J.W., Milnes, J.T., El-Haou, S., Jackson, C., Gkatzis, K., Elliott, D.A., Chuva de Sousa Lopes, S.M., Mummery, C.L., et al. (2015). Atrial-like cardiomyocytes from human pluripotent stem cells are a robust preclinical model for assessing atrial-selective pharmacology. EMBO Mol Med 7, 394–410.

Francou, A., and Kelly, R.G. (2016). Properties of Cardiac Progenitor Cells in the Second Heart Field. In Etiology and Morphogenesis of Congenital Heart Disease: From Gene Function and Cellular Interaction to Morphology, T. Nakanishi, R.R. Markwald, H.S. Baldwin, B.B. Keller, D. Srivastava, and H. Yamagishi, eds. (Tokyo: Springer), p.

Garcia-Martinez, V., and Schoenwolf, G.C. (1993). Primitive-Streak Origin of the Cardiovascular System in Avian Embryos. Developmental Biology 159, 706–719.

Gavrilov, S., and Lacy, E. (2013). Genetic dissection of ventral folding morphogenesis in mouse: embryonic visceral endoderm-supplied BMP2 positions head and heart. Curr Opin Genet Dev 23, 461–469.

Gessert, S., and Kühl, M. (2010). The Multiple Phases and Faces of Wnt Signaling During Cardiac Differentiation and Development. Circulation Research 107, 186–199.

High, F.A., Jain, R., Stoller, J.Z., Antonucci, N.B., Lu, M.M., Loomes, K.M., Kaestner, K.H., Pear, W.S., and Epstein, J.A. (2009). Murine Jagged1/Notch signaling in the second heart field orchestrates Fgf8 expression and tissue-tissue interactions during outflow tract development. J Clin Invest 119, 1986–1996.

Hochgreb, T., Linhares, V.L., Menezes, D.C., Sampaio, A.C., Yan, C.Y.I., Cardoso, W.V., Rosenthal, N., and Xavier-Neto, J. (2003). A caudorostral wave of RALDH2 conveys anteroposterior information to the cardiac field. Development 130, 5363–5374.

Ivanovitch, K., Soro-Barrio, P., Chakravarty, P., Jones, R.A., Bell, D.M., Mousavy Gharavy, S.N., Stamataki, D., Delile, J., Smith, J.C., and Briscoe, J. (2021). Ventricular, atrial, and outflow tract heart progenitors arise from spatially and molecularly distinct regions of the primitive streak. PLoS Biol 19, e3001200.

Jain, R., Li, D., Gupta, M., Manderfield, L.J., Ifkovits, J.L., Wang, Q., Liu, F., Liu, Y., Poleshko, A., Padmanabhan, A., et al. (2015). HEART DEVELOPMENT. Integration of Bmp and Wnt signaling by Hopx specifies commitment of cardiomyoblasts. Science 348, aaa6071.

Keegan, B.R., Meyer, D., and Yelon, D. (2004). Organization of cardiac chamber progenitors in the zebrafish blastula. Development 131, 3081–3091.

Kelly, R.G., Buckingham, M.E., and Moorman, A.F. (2014). Heart fields and cardiac morphogenesis. Cold Spring Harb Perspect Med 4, a015750.

Kloesel, B., DiNardo, J.A., and Body, S.C. (2016). Cardiac Embryology and Molecular Mechanisms of Congenital Heart Disease – A Primer for Anesthesiologists. Anesth Analg 123, 551–569.

Lebensohn, A.M., and Rohatgi, R. (2018). R-spondins can potentiate WNT signaling without LGRs. ELife 7, e33126.

Lee, J.H., Protze, S.I., Laksman, Z., Backx, P.H., and Keller, G.M. (2017). Human Pluripotent Stem Cell-Derived Atrial and Ventricular Cardiomyocytes Develop from Distinct Mesoderm Populations. Cell Stem Cell 21, 179–194.e4.

Lescroart, F., Chabab, S., Lin, X., Rulands, S., Paulissen, C., Rodolosse, A., Auer, H., Achouri, Y., Dubois, C., Bondue, A., et al. (2014). Early lineage restriction in temporally distinct populations of Mesp1 progenitors during mammalian heart development. Nat Cell Biol 16, 829–840.

Lescroart, F., Wang, X., Lin, X., Swedlund, B., Gargouri, S., Sànchez-Dànes, A., Moignard, V., Dubois, C., Paulissen, C., Kinston, S., et al. (2018). Defining the earliest step of cardiovascular lineage segregation by single-cell RNA-seq. Science 359, 1177–1181.

Li, G., Xu, A., Sim, S., Priest, J.R., Tian, X., Khan, T., Quertermous, T., Zhou, B., Tsao, P.S., Quake, S.R., et al. (2016). Transcriptomic Profiling Maps Anatomically Patterned Subpopulations among Single Embryonic Cardiac Cells. Dev Cell 39, 491–507.

Lin, S.-C., Dollé, P., Ryckebüsch, L., Noseda, M., Zaffran, S., Schneider, M.D., and Niederreither, K. (2010). Endogenous retinoic acid regulates cardiac progenitor differentiation. Proc Natl Acad Sci U S A 107, 9234–9239.

Mantri, M., Scuderi, G.J., Abedini-Nassab, R., Wang, M.F.Z., McKellar, D., Shi, H., Grodner, B., Butcher, J.T., and De Vlaminck, I. (2021). Spatiotemporal single-cell RNA sequencing of developing chicken hearts identifies interplay between cellular differentiation and morphogenesis. Nat Commun 12, 1771.

Meilhac, S.M., Lescroart, F., Blanpain, C., and Buckingham, M.E. (2014). Cardiac Cell Lineages that Form the Heart. Cold Spring Harb Perspect Med 4, a013888.

Miquerol, L., and Kelly, R.G. (2013). Organogenesis of the vertebrate heart. Wiley Interdiscip Rev Dev Biol 2, 17–29.

Niderla-BieliŃska, J., Jankowska-Steifer, E., Flaht-Zabost, A., Gula, G., Czarnowska, E., and Ratajska, A. (2019). Proepicardium: Current Understanding of its Structure, Induction, and Fate. The Anatomical Record 302, 893–903.

Niederreither, K., Vermot, J., Messaddeq, N., Schuhbaur, B., Chambon, P., and Dollé, P. (2001). Embryonic retinoic acid synthesis is essential for heart morphogenesis in the mouse. Development 128, 1019–1031.

Ola, R., Lefebvre, S., Braunewell, K.-H., Sainio, K., and Sariola, H. (2012). The expression of Visinin-like 1 during mouse embryonic development. Gene Expr Patterns 12, 53–62.

Perl, E., and Waxman, J.S. (2019). Reiterative Mechanisms of Retinoic Acid Signaling during Vertebrate Heart Development. J Dev Biol 7, E11.

Perl, E., and Waxman, J.S. (2020). Retinoic Acid Signaling and Heart Development. Subcell Biochem 95, 119–149.

Pierpont, M.E., Brueckner, M., Chung, W.K., Garg, V., Lacro, R.V., McGuire, A.L., Mital, S., Priest, J.R., Pu, W.T., Roberts, A., et al. (2018). Genetic Basis for Congenital Heart Disease: Revisited: A Scientific Statement From the American Heart Association. Circulation 138, e653–e711.

Piersma, A.H., Hessel, E.V., and Staal, Y.C. (2017). Retinoic acid in developmental toxicology: Teratogen, morphogen and biomarker. Reprod Toxicol 72, 53–61.

Pradhan, A., Zeng, X.-X.I., Sidhwani, P., Marques, S.R., George, V., Targoff, K.L., Chi, N.C., and Yelon, D. (2017). FGF signaling enforces cardiac chamber identity in the developing ventricle. Development 144, 1328–1338.

Rana, M.S., Théveniau-Ruissy, M., De Bono, C., Mesbah, K., Francou, A., Rammah, M., Domínguez, J.N., Roux, M., Laforest, B., Anderson, R.H., et al. (2014). Tbx1 Coordinates Addition of Posterior Second Heart Field Progenitor Cells to the Arterial and Venous Poles of the Heart. Circulation Research 115, 790–799.

Roux, M., Laforest, B., Capecchi, M., Bertrand, N., and Zaffran, S. (2015). Hoxb1 regulates proliferation and differentiation of second heart field progenitors in pharyngeal mesoderm and genetically interacts with Hoxa1 during cardiac outflow tract development. Dev Biol 406, 247–258.

de Soysa, T.Y., Ranade, S.S., Okawa, S., Ravichandran, S., Huang, Y., Salunga, H.T., Schricker, A., Del Sol, A., Gifford, C.A., and Srivastava, D. (2019). Single-cell analysis of cardiogenesis reveals basis for organ-level developmental defects. Nature 572, 120–124.

Stefanovic, S., and Zaffran, S. (2017). Mechanisms of retinoic acid signaling during cardiogenesis. Mech Dev 143, 9–19.

Tyser, R.C.V., Ibarra-Soria, X., McDole, K., Arcot Jayaram, S., Godwin, J., van den Brand, T.A.H., Miranda, A.M.A., Scialdone, A., Keller, P.J., Marioni, J.C., et al. (2021). Characterization of a common progenitor pool of the epicardium and myocardium. Science 371, eabb2986.

Welsh, I.C., Hart, J., Brown, J.M., Hansen, K., Rocha Marques, M., Aho, R.J., Grishina, I., Hurtado, R., Herzlinger, D., Ferretti, E., et al. (2018). Pbx loss in cranial neural crest, unlike in epithelium, results in cleft palate only and a broader midface. J Anat 233, 222–242.

Xavier-Neto, J., Neville, C.M., Shapiro, M.D., Houghton, L., Wang, G.F., Nikovits, W., Stockdale, F.E., and Rosenthal, N. (1999). A retinoic acid-inducible transgenic marker of sino-atrial development in the mouse heart. Development 126, 2677–2687.

Yutzey, K.E., and Bader, D. (1995). Diversification of Cardiomyogenic Cell Lineages During Early Heart Development. Circulation Research 77, 216–219.

Zaffran, S., Kelly, R.G., Meilhac, S.M., Buckingham, M.E., and Brown, N.A. (2004). Right Ventricular Myocardium Derives From the Anterior Heart Field. Circulation Research 95, 261–268.

Zamir, L., Singh, R., Nathan, E., Patrick, R., Yifa, O., Yahalom-Ronen, Y., Arraf, A.A., Schultheiss, T.M., Suo, S., Han, J.-D.J., et al. Nkx2.5 marks angioblasts that contribute to hemogenic endothelium of the endocardium and dorsal aorta. ELife 6, e20994.

Zhang, Q., Jiang, J., Han, P., Yuan, Q., Zhang, J., Zhang, X., Xu, Y., Cao, H., Meng, Q., Chen, L., et al. (2011). Direct differentiation of atrial and ventricular myocytes from human embryonic stem cells by alternating retinoid signals. Cell Res 21, 579–587.

